# A transcriptome map reveals conserved and lineage-specific sRNAs that regulate motility and metabolism across *Vibrio*

**DOI:** 10.64898/2026.08.28.747715

**Authors:** Zhizhou Jia, Han Zhang, Daniel Falush, Yanjie Chao, Sarah L. Svensson

## Abstract

Bacterial sRNAs are pervasive post-transcriptional regulators, yet how they arise, evolve, and decay remains poorly understood. Here, we provide a high-resolution transcriptome map and curated sRNA set for the pathogen *Vibrio parahaemolyticus*. We identify over 100 sRNAs, including broadly conserved, lineage-specific, and previously unidentified transcripts, as well as dual-function regulatory/coding sRNAs. Functional analysis of several examples highlights conserved and lineage-specific regulators of metabolism and flagella. Broadly conserved VcrX represses chitin utilization genes and may regulate *Vibrio* Spot 42, which we confirm is translated. We expand on FlaX regulation of polar flagella across the genus by demonstrating that the sRNA differentially activates/represses downstream flagellins, with a potential FlaX sponge mediating feedback in specific clades. We further show that *V. parahaemolyticus*, but not *V. cholerae*, RyhB is translated into a Cys-rich small protein that could regulate related pathways. Together, these findings establish a resource for *Vibrio* and a platform for comparative studies of post-transcriptional regulation, enabling investigation of how sRNAs and their regulatory networks evolve.

## Introduction

Bacterial small RNAs (sRNAs) are widespread post-transcriptional regulators that control phenotypes ranging from metabolism and motility to host colonization and virulence. Most act through short base-pairing interactions with target mRNAs, typically interacting with 5’ UTRs (untranslated regions) to repress translation. However, sRNAs can also regulate mRNA stability or activate translation, and can also bind mRNA coding regions, other sRNAs, or proteins^1^. A handful of “dual-function” sRNAs encode small proteins (<50 amino acids) that regulate targets in the related pathways^2^. This diversity in function allows sRNAs to provide a variety of distinct regulatory roles from that of transcription factors, ranging from control of broad regulons to fine-tuning of individual targets within network motifs. In Gammaproteobacteria, expression or base-pairing activity of many sRNAs is also organized by RNA chaperones such as Hfq and ProQ.

How sRNAs arise, evolve, and decay remains poorly understood^3–5^. Compared with transcription factors and other protein regulators, sRNAs provide a distinct evolutionary substrate, with fewer constraints on their sequence. Selection on an sRNA also depends on the expression and dynamics of target mRNAs. Riboregulators tend to be younger in age and are often more lineage specific: only ∼60% of sRNA families are conserved between species, compared with ∼90% of protein regulators^3^, and most have arisen within species or even strains^4,6^. New sRNAs can more easily arise *de novo*, for example via single nucleotide polymorphisms (SNPs) that generate a new promoter or cleavage site, or as the result of genome rearrangements^5^. RNAs are also found on dynamic, horizontally transferred elements such as prophages and pathogenicity islands, where cross regulation between the core and acquired genome evolves^7^. Functional homologs, such as a variety of sRNAs regulating iron metabolism, may also evolve convergently without detectable sequence similarity.

These characteristics, as well as the short length of sRNAs and their lack of the well-defined domains of protein regulators add up to a challenging landscape for comparative studies, although a handful are available, especially in the Enterobacteriaceae ^8–10^ but also in other clades^11–13^. The recent expansion of the known sRNA universe has been driven by annotation-independent RNA-seq-based technologies, for example that map transcription start sites or RNA chaperone interactomes, which have revealed functional sRNAs encoded in diverse genomic contexts^14,15^. Applying these methods across a taxon, especially one that is well characterized and genetically tractable, might complement comparative genomics of sRNAs, which can be challenging^16–18^.

The *Vibrio* genus and related Vibrionaceae encompass free-living, symbiotic, as well as pathogenic species, such as *Vibrio cholerae, V. vulnificus*, *V. parahaemolyticus*. Several species are models for studying, e.g., symbiosis, quorum sensing, motility, and bioengineering. Vibrios are well suited for evolutionary studies due to their ecological and genomic diversity, high rates of recombination and horizontal gene transfer, known genotype-phenotype-ecology links, as well as the availability of genome sequences and genetically tractable strains. Their tendency to generate pathogenic lineages and human/climate-driven changes in their distribution also makes understanding *Vibrio* adaptation crucial for global public health^19^. Recently, *V. cholerae* and its phages have also emerged as models for studying sRNA-based regulation (reviewed in^20^), including strain-specific sRNAs^21^. However, it is unclear how these insights apply across the genus.

Here, we provide high-resolution RNA-seq datasets and a curated set of candidate sRNAs for *V. parahaemolyticus*, filling a gap in foundational resources for *Vibrio* and enabling comparative analysis of sRNAs across this well-studied bacterial genus. *V. parahaemolyticus* is endemic to coastal marine/estuarine environments and a leading seafood-borne and aquaculture pathogen worldwide. Its extensive diversity, distinct geographic populations, and exceptionally high recombination rate make it a powerful model for linking genotype to phenotype and ecology^22^. We curate sRNAs based on their expression and conservation, revealing both conserved and lineage-specific transcripts. Functional analysis of selected examples highlights shared and divergent mechanisms regulating motility and metabolism across the genus, including potential new modes of post-transcriptional regulation. Together with extensive resources and insight from *V. cholerae* and other Gammaproteobacteria, this establishes *Vibrio* as a powerful system for investigating the evolution of sRNAs and post-transcriptional regulatory networks.

## Results

### High resolution transcriptome datasets for *Vibrio parahaemolyticus*

To provide transcriptome coordinates and expression information for *V. parahaemolyticus*, we generated four different types of RNA-seq dataset (**Fig. 1A**): conventional RNA-seq, differential RNA-seq (dRNA-seq) and Cappable-seq for RNA 5’ ends and transcription start sites (TSS) ^23,24^, as well as term-seq for 3’ ends ^27^. We used the available wild-type strain VN-5003-Wue (NZ_CM177759.1), but reads were mapped to the widely used reference strain RIMD 2210633, a member of the pandemic clone lineage ^28^. The percentage of mapped reads was high (>95%) despite the data being generated in a different strain (**Table S1**). The datasets are exemplified by coverage at the *vqmRA* locus, encoding a quorum sensing-related sRNA and its regulator ^29^ (**Fig. 1B; Fig. S1A**). Processed coverage files, along with an updated annotation for strain RIMD 2210633, are available at https://doi.org/10.6084/m9.figshare.32682531.

**Figure 1.**
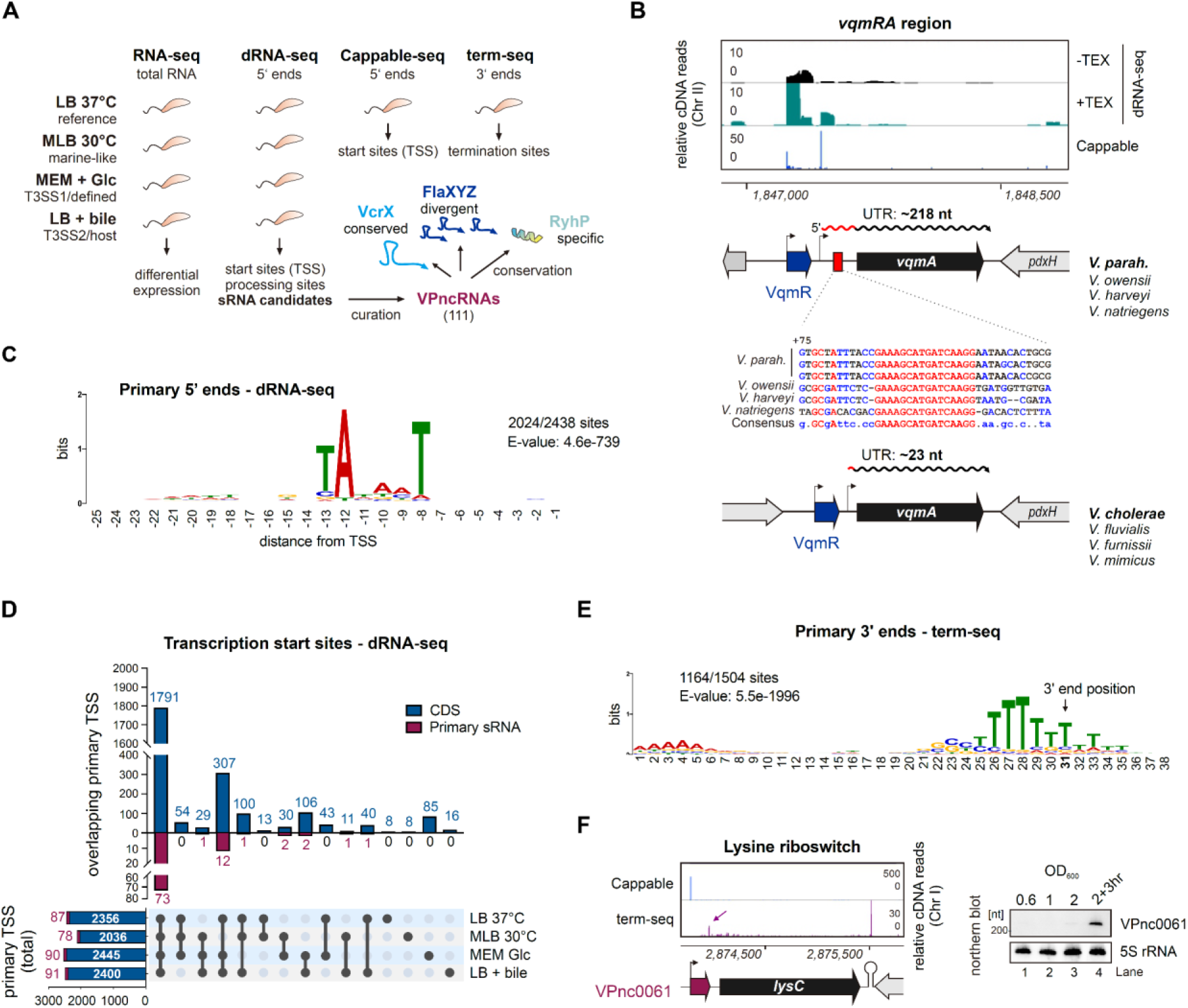
*V. parahaemolyticus* transcriptome datasets map and characterize RNA 5’ and 3’ ends. **(A)** Schematic of generated RNA-seq datasets, downstream analysis, and insights. **(B)** Expression and conservation of the *vqmRA* locus in *Vibrio*. *Top*: cDNA library coverage for 5’ ends. *Bottom*: Schematic of genomic organization/5’ UTR length for two types of *vqmRA* ^25^ (**Fig. S2B**) and conservation of a 16 bp motif in the long 5’ UTR of *V. parahaemolyticus* and related species. **(C)** MEME motif analysis of the 50 bp upstream of primary TSS identified by dRNA-seq in LB. While 50 bp regions were used for prediction, positions < -25 nt are omitted for space. **(D)** Upset plot of TSS identified in four growth conditions by dRNA-seq. **(E)** Intrinsic terminator motif identified in term-seq mapped 3’ ends +/- 30 nt by MEME ^26^. The arrow indicates the middle nucleotide of the 61 bp region used for prediction, positions >38 nt are omitted for space. **(F)** (*Left*) cDNA library coverage at *lysC.* Arrow: 3’-end of termination product (VPnc0061). (*Right*) VPnc0061 northern blot for RNA from bacteria growing in MLB. 5S rRNA: loading control.

For conventional RNA-seq (**Table S2**), bacteria were grown in four different conditions established previously: LB37 (LB at 37°C), MLB30 (LB + 3% NaCl at 30°C), T3SS1-inducing (Dulbecco’s Modified Eagle Medium, DMEM at 37°C), and T3SS2-inducing (LB + 0.05% bile salts at 37°C) ^30^, with all harvested in mid- to late-log phase. Genes encoding Type III secretion system 1 (T3SS1) were more highly expressed in the T3SS1 condition (vs. LB at 37°C), and glucose transport genes were upregulated, consistent with growth in DMEM (**Fig. S1B, C)**. While we could not confirm induction of T3SS2 (the system is absent from VN-5003-Wue), two efflux pumps were induced in T3SS2 libraries vs. LB, consistent with bile salt treatment (**Fig. S1D, E**).

To increase the number of TSS detected, we also generated dRNA-seq data under the four different conditions above. Both VqmR and *vqmA* had clear primary TSS, based on enrichment in TEX (terminator exonuclease) libraries vs. mock-treated controls (**Fig. 1B**). Analyzing data from all four conditions using ANNOgesic ^31^ mapped 10,178 TSS (**Table S3**). We subsequently re-assigned each TSS as primary, secondary, internal, or antisense using our updated annotation with 111 VPncRNAs (see following section and **Table S4**). MEME motif analysis of the 50 nt upstream of each primary TSS detected in LB revealed a canonical -10 box sequence located ∼8-15 nt upstream (**Fig. 1C**). The list of primary TSS included 2,692 associated with the 4,831 annotated ORFs in strain RIMD 2210633 (NCBI annotation, 05/30/2021) and 93 associated with VPncRNAs (**Fig. 1D; Table S3**). Only 1,864 primary TSS were detected in all four conditions (69.2%), demonstrating the utility of assaying different media.

The TSS map confirmed previously suggested regulatory diversity in *Vibrio*. We noticed that the *vqmA* TSS in *V. parahaemolyticus* generates a much longer 5’ UTR (>200 nt) than in *V. cholerae* (∼25 nt) (**Fig. 1B**) ^29^. Closer inspection revealed that organization at the *vqmRA* locus falls into two main types in *Vibrio* species (**Fig. S2**), as was reported before based on putative *V. parahaemolyticus* and *V. cholerae* promoter sequences ^25^. Alignments of the longer *vqmA* 5’ UTR revealed a striking 16 bp conserved sequence that could mediate post-transcriptional regulation (**Fig. 1B**) and might also be responsible for differential regulation ^25^.

Based on the TEX sensitivity, rather than resistance, of 5’ monophosphates, dRNA-seq can also identify processing sites. For example, the monophosphorylated, mature 5’ end of RNase E-processed MicX ^32,33^ was enriched in -TEX libraries, while the TSS driving transcription of the 5’-triphosphorylated precursor was enriched in +TEX libraries (**Fig. S3A**). Processing sites were detected with ANNOgesic using parameters optimized with manually curated sites, followed by inspection of coverage, resulting in a set of 69 putative cleavage sites (**Table S5**). These positions included several associated with known or uncharacterized sRNAs encoded in 3’ UTRs, indicating they might be generated by processing. Some processing sites were detected in only a single condition (**Fig. S3B**), such as for 3’ UTR-derived VPnc0074 in the T3SS2 library (**Fig. S3C**), although this could reflect expression rather than processing differences.

We also generated Cappable-seq libraries, based on specific capping of triphosphorylated 5’ ends with Vaccinia capping enzyme ^24^, in the LB condition for comparison with dRNA-seq (**Table S6**). Cappable-seq peaks were consistent with dRNA-seq TSS by visual inspection, and MEME analysis identified a similar -10 motif upstream of primary TSS as for dRNA-seq (**Fig. 1C & S3D**). However, automated detection revealed that only 1288 primary TSS overlapped between the methods (**Fig. S3E,** *left*). This might reflect analysis methods, etc., but suggests that each approach has distinct specificity or sensitivity issues. To explore this, we looked for an overlap between Cappable-seq TSS and dRNA-seq processing sites. Although peak height was generally low, we found 16 Cappable-seq TSS peaks overlapped with dRNA-seq processing sites (**Fig. S3E**, *right*). This included three associated with sRNAs that are processed in *V. cholerae* (MicX, FarS, FlaX; **Fig. S3A, F, G**) ^32,34,35^. Peaks overlapping cleavage sites tended to have higher expression in total RNA-seq (**Fig. S3H**) and may reflect carry-over during library preparation.

Finally, we mapped transcript 3’ ends using term-seq. Using a previous method ^36^, we identified 5,236 3’ ends, including 1,504 “primary” 3’ ends within 100 bp downstream of annotated stop codons (**Table S7**). MEME motif analysis ^26^ of the 30 nt surrounding primary 3’ ends identified a poly-T hallmark of Rho-independent terminators in 1,164 out of 1,504 sites (**Fig. 1E**). The list of 3’ ends included those associated with known regulatory elements, such as a lysine-sensitive riboswitch in *lysC*/VP2715 (**Fig. 1F**) ^37^. For some longer operon ends, term-seq peaks were not strong, such as at the end of the VPA1197-VPA1201 nitrate reductase operon (**Fig. S4A**). This region caught our attention because of a putative >200 bp “empty” 3’ UTR, which encodes a small, translated 44 amino acid protein specific to *Vibrio* (TIGR02808) that may be a functional homolog of NapE and interact with NapC, encoded immediately upstream (**Fig. S4B, C**). Nonetheless, our dataset should allow for mapping 3’ UTR-derived sRNAs and 3’ ends resulting from control of transcription elongation and will complement the *V. cholerae* 3’ end map ^33^.

### A repertoire of conserved and variable sRNAs

While sRNAs have been characterized in *V. cholerae* ^20^, it is not clear how many are conserved in *Vibrio* or if species-specific examples exist. After manually inspecting ANNOgesic predictions for genomic context and coverage to remove, *e.g*., those overlapping tRNAs or rRNAs, we obtained 101 candidates (**Fig. 2**; **Table S4**). These included housekeeping RNAs (SRP [signal recognition particle] RNA, RnpB, tmRNA), Gammaproteobacterial sRNAs (Spot 42, GcvB, RyhB), and most characterized *V. cholerae* sRNAs, such as VrrA, VadR, VcdR, VqmR, MicX, FarS, MltS, FlaX, and MicV ^29,32,34,35,38–42^. We added another ten via inspection of conserved genomic regions or BLAST with *V. cholerae* homologs, such as TfoR, CarZ, OppX, and VSsrna24, as well as potential homologs of the flagellin 3’ UTR-derived sRNAs Vcr77 and Vcr78 ^33,35,43^ that were missed by ANNOgesic, possibly due to low expression.

**Figure 2.**
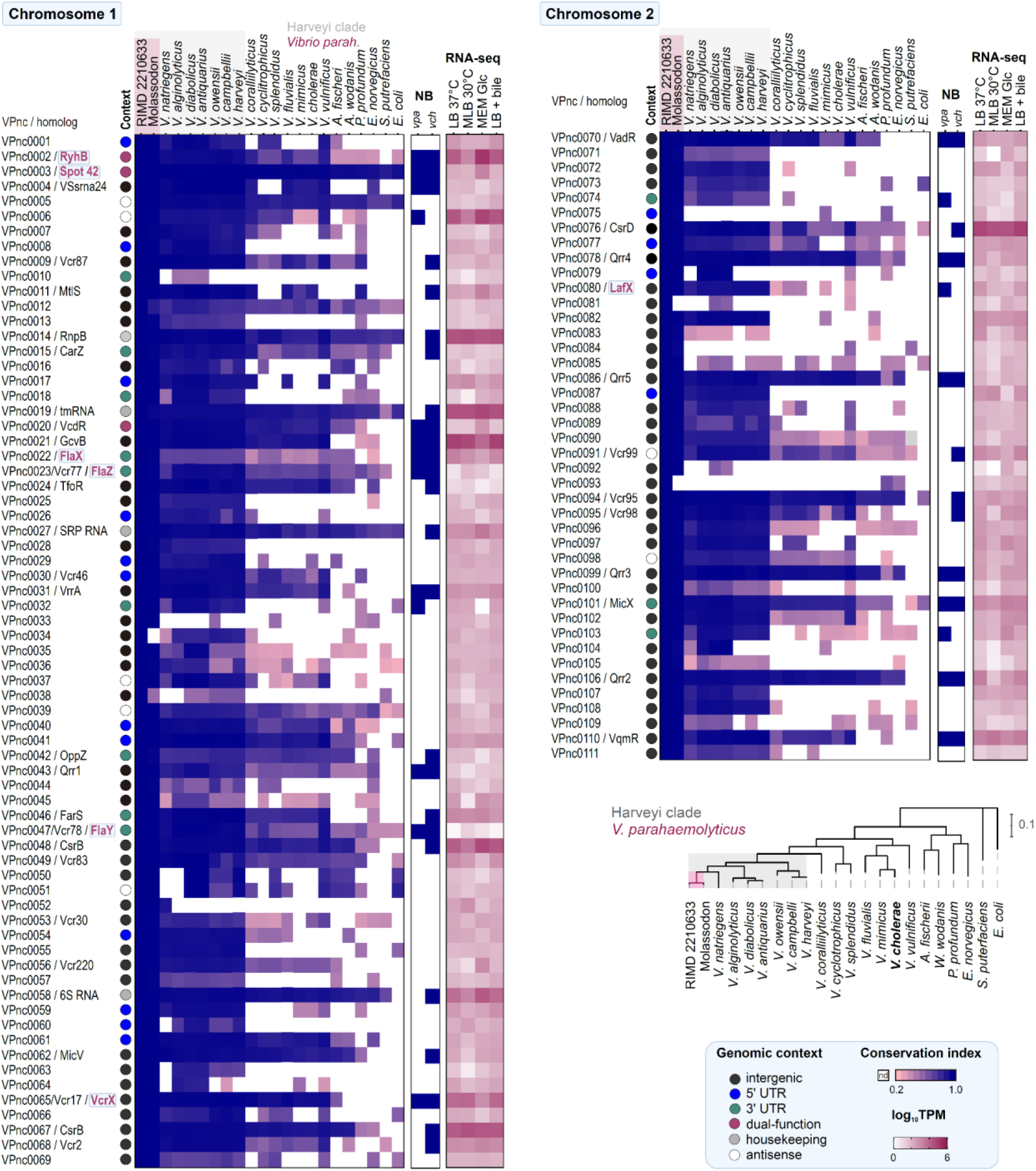
Genomic context, conservation, and expression for *V. parahaemolyticus* non-coding RNA (VPnc) candidates on two chromosomes of strain RIMD 2210633. See also **Table S4**. Common names are based on homology. Candidates named in pink were studied in this work. Those validated by northern blot in this study (*vpa*, **Fig. S6**) or previous work (*vch*) are also indicated. RNA-seq: mean log_10_TPM (transcripts per million) in the indicated conditions, n=2. Conservation index: square root of the product of % identity and coverage. nd: not detected (conservation index <0.2, percent ID <50, e-value >1) based on BLASTn. *Bottom right*: Maximum likelihood phylogenetic tree based on *gyrB*. The consensus tree of 100 bootstrappings is shown.

We named these 111 candidates VPnc0001-VPnc0111 in order of genome position (**Fig. 2**; **Table S4**). As in other bacteria, the VPncRNAs are encoded in diverse genomic contexts, with 12 in 3’ UTRs. Some of these appear to be specific to our study and potentially to *V. parahaemolyticus*, supporting the idea that 3’ UTRs are a “playground” for sRNA evolution ^5,44^. Those with established functions tended to be more conserved and like housekeeping genes, encoded on Chromosome 1, as well as more highly expressed. The sRNAs encoded on Chromosome 1 also tended to be more conserved than those on Chromosome 2, in line with previous observations in *V. splendidus* ^45^.

To confirm which of the 111 VPncRNAs are shared with *V. cholerae*, we performed BLAST searches against the *V. cholerae* C6706 genome. This identified matches for only 63 (**Fig. S5A; Table S4**) (E-value <0.1, coverage & percent identity >50%): four housekeeping RNAs, 23 previously studied sRNAs, ten with homology to unstudied *V. cholerae* sRNAs (Vcrs), and 26 that mapped elsewhere in the genome. The remaining 37 VPncRNAs may be restricted to *V. parahaemolyticus* and its relatives. We also performed the reverse search, BLASTing 158 RNAs from *V. cholerae* identified by dRNA-seq and Hfq RIP-seq ^29,34^ against the *V. parahaemolyticus* RIMD 2210633 genome. This recovered 97 matches (E-value <0.1, coverage/identity >50%; **Table S8**), including 4/4 housekeeping RNAs and 60/154 sRNAs. Of the sRNAs, 35 overlapped with the 111 VPncRNAs (**Fig. S5B**). Most of the remaining 25 matches mapped within ORFs or intergenic regions with limited cDNA coverage, making it unclear if they are *bona fide* sRNAs in *V. parahaemolyticus*. While several corresponded to 5′/3′ UTRs, it was not clear from RNA-seq coverage that these are independent transcripts (**Fig. S5C–F**). Northern analysis is needed to determine if they are sRNAs in *V. parahaemolyticus,* or if they represent species-specific sRNAs.

There were several notable absences of studied *V. cholerae* sRNAs, including CoaR and sponge sRNA QrrX ^46,47^, which we also could not find even by manual inspection of adjacent homologous genes. Also absent were the ToxT-regulated sRNAs TarA/TarB, which are encoded on the *Vibrio* pathogenicity island VPI-1 ^48^. We also did not recover an sRNA in the *V. parahaemolyticus ompU* (VP2467) 3’ UTR, which in *V. cholerae* encodes recently described OueS (**Fig. S5G**) ^21^. Overall, our inspection suggests that the *Vibrio* genus encodes both highly conserved and lineage specific sRNAs. However, it is also possible that they are not expressed under the conditions we assayed and/or have diverged beyond recognition by routine homology searches.

To complement RNA-seq data (**Fig. 2**; **Table S2, S4**), we confirmed the expression of several sRNAs using northern blot analysis with RNA from the wild-type strain RIMD 2210633 harvested at different growth phases in rich media, as well as isogenic Δ*hfq* and Δ*proQ* mutants sampled in log phase (**Fig. S6A-D**). We confirmed homologs of several known sRNAs, including broadly conserved Gammaproteobacterial sRNAs (Spot 42, RyhB, GcvB; **Fig. S6A**) as well as several studied *V. cholerae* sRNAs (**Fig. S6B**), with many showing differential expression between growth phases. For example, Spot 42 levels decreased at later growth phases, while quorum sensing related VqmR increased on entry into stationary phase.

We also confirmed the expression of uncharacterized 3’ UTR-derived and intergenic sRNAs (**Fig. S6C, D**). VPnc0103, a putative homolog of Vcr106, is encoded downstream of an ABC transporter, accumulates in stationary phase and might be expressed as a precursor and processed (**Fig. S6C, E**) ^29,33^. To our knowledge, expression of the *V. cholerae* homolog Vcr106 was not previously confirmed. The other two sRNAs, VPnc0074 (**Fig. S3C**) and VPnc0032 (**Fig. S6F**) appear to be absent from *V. cholerae* and restricted to the Harveyi clade (**Fig. 2**). VPnc0074 is encoded in the 3’ UTR of the *ompW* porin (VPA0096) and is induced in response to bile salts (**Fig. S3C**). VPnc0032 is encoded in the 3’ UTR of a dicarboxylate transporter and repressed, along with its parental mRNA, in the presence of glucose (**Fig. S6F**). Finally, we also confirmed several uncharacterized intergenic sRNAs (**Fig. S6D**). This included a >300 nt transcript, VPnc0006 (**Fig. S6G**), which is highly abundant in RNA-seq data and encoded antisense to VP0111 (hypothetical protein), VPnc0035 (**Fig. S6H**), downstream of *fdhD*, and conserved VPnc0065 (VcrX, next section), which accumulates stationary phase (**Fig. 2**; **Fig. S6D**).

*Vibrio* species encode Hfq and ProQ RNA chaperones. While several sRNAs (RyhB, Spot 42, VadR, MicX, VPnc0074, VPnc0006, VPnc0035, and VPnc0065/VcrX) showed *hfq*-dependent expression in log phase, many sRNAs that are known to be functional or Hfq-associated in *V. cholerae* were not strongly affected in Δ*hfq* (**Fig. S6A-D**), such as VqmR ^29,34^, or were even upregulated (*e.g*., VrrA, **Fig. S6B,** lanes 2 vs. 6). These effects may be due to pleiotropic transcriptome changes or growth differences in Δ*hfq,* as exemplified by upregulation of the density-dependent Qrr RNAs. ProQ absence also did not affect many of the sRNAs we tested in log phase, other than mildly lower levels of FlaX (**Fig. S6B,** lanes 2 vs. 7), which binds ProQ in *V. cholerae* ^35^. The reason for the independence of many *Vibrio* sRNAs from Hfq and ProQ for expression is unclear. However, *Vibrio* Hfq is markedly different from that of enterobacteria ^49^ and might have a distinct cellular role. The chaperones might still affect sRNA function.

### The conserved *Vibrio* sRNA VcrX regulates chitin usage

The most conspicuous sRNA that we annotated was VPnc0065 (**Fig. 3A**), which we rename <u>Vcr</u>X (*Vibrio* carbon/chitin regulator) based on analyses described below. VcrX (Vcr17 in *V. cholerae*) was as conserved and abundantly expressed as other well-studied *Vibrio* sRNAs (**Fig. 2**; **Fig. 3B**), strongly suggesting it has a central function in the genus . However, the sRNA is still uncharacterized, although several targets were detected and confirmed for Vcr17 as part of an Hfq RIL-seq screen ^46^.

**Figure 3.**
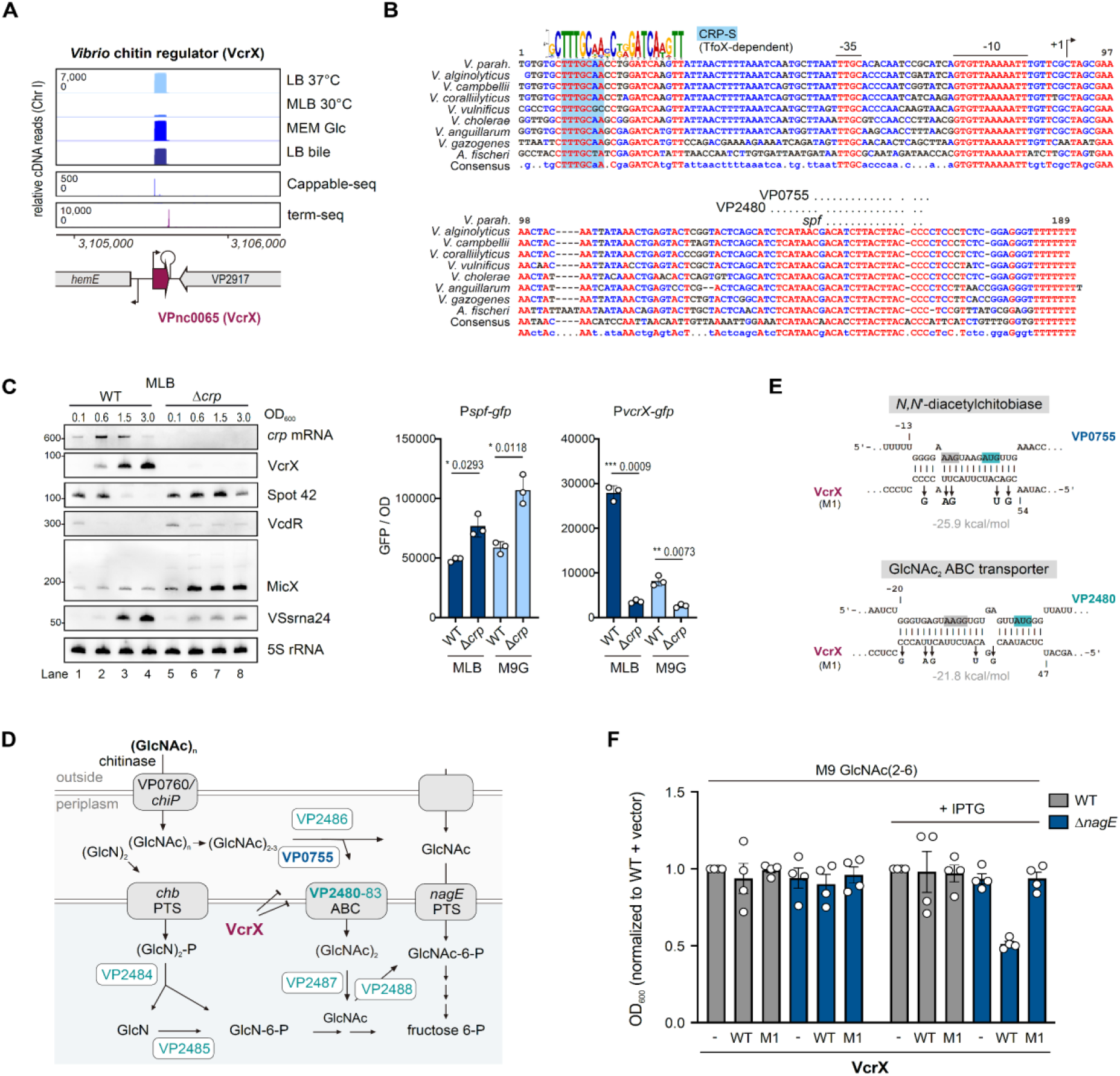
The conserved *Vibrio* sRNA VcrX regulates chitosugar usage. **(A)** cDNA coverage and context for *V. parahaemolyticus* VcrX. **(B)** Alignment of *vcrX* regions identified in *Vibrio* species and *Aliivibrio fischeri*. Bent arrow: TSS. Sequence motif: MEME ^26^ e-value 8e-44). Blue boxes: putative CRP-S (TfoX-dependent, based on ^51^) motif. Horizontal dotted lines: VP2480/VP0755 seeds and putative Spot 42 interaction region in *V. parahaemolyticus*. **(C)** (*Left*) Northern blot of CRP- or carbon source-related sRNAs in WT/Δ*crp* strains at different optical densities in MLB. OD_600_ 0.1: early log phase, 0.6: log, 1.5: early stationary, 3.0: stationary. 5S rRNA: loading control. Representative of two independent experiments. (*Right*) *vcrX* and *spf* GFP transcriptional fusion expression in WT/Δ*crp* in stationary phase in MLB. GFP fluorescence was normalized to culture OD_600_. n=3, Student’s *t*-test. *p<0.05, **p<0.01, ***p<0.001. **(D)** Predicted base-pairing for VcrX and VP2480/VP0755. Grey: RBS. Green: start codons. M1: point mutations. mRNA numbering: vs. start codon. **(E)** Schematic of *V. parahaemolyticus* chitin utilization ^53,55^. Bold targets: direct. Green: VP2480 operon. PTS: phosphotransferase system. ABC: ATP synthase-binding cassette transporter. **(F)** Growth of strains carrying an empty vector or VcrX WT/M1 with GlcNAc(2-6) as carbon source. *nagE*: GlcNAc transporter. n=3 independent cultures. Mean vs. WT + empty vector without IPTG.

Northern blot analysis confirmed that *V. parahaemolyticus* VcrX is a 95 nt, Hfq-dependent sRNA that accumulates upon entry into stationary phase in rich media (**Fig. S6D**), as was observed in *V. cholerae* ^29^. VcrX was also repressed in the MEM + glucose condition, and we noticed that VcrX expression tended to be inversely related to Spot 42 levels across growth phases in rich media and with various carbon sources (**Fig. 2, S6A, D; Fig. S7A**), which was higher at early growth phases in rich media and de-repressed by glucose. Spot 42 regulates carbon metabolism in diverse Gammaproteobacteria, suggesting that VcrX could regulate genes in related pathways.

Spot 42 is repressed by CRP (cAMP receptor protein) binding to its promoter in the absence of glucose, when cAMP levels are high. We identified a highly conserved motif upstream of the VcrX promoter -35 region (CTTTGCnnnCnnGATCA; **Fig. 3B**). This consensus does not match the canonical CRP site. Nonetheless, in line with CRP regulation, VcrX expression was absent in a Δ*crp* mutant, while expression of Spot 42 was increased in the same strain (**Fig. 3C**, *left*). Several other sRNAs previously linked to carbon source regulation were also dysregulated in the CRP mutant but did not show the striking inverse correlation with Spot 42. We observed the same trend for promoter activity (**Fig. 3C**, *right*; **Fig. S7A**). While it remains to be seen if this is via direct CRP binding, a possibility is that the motif is a degenerate CRP-S site, and requires the chitin-dependent co-activator TfoX^50,51^, unlike the canonical CRP-N site of Spot 42 ^50,51,50,51^ Regulation in response to different carbon sources and by CRP strongly suggests that VcrX is part of the carbon source regulatory scheme of *Vibrio*.

To investigate how VcrX might participate in carbon source regulation, we predicted mRNA targets using CopraRNA ^52^ and homologs from several species (*V. parahaemolyticus*, *V. campbellii*, *V. harveyi*, *V. vulnificus*, and *V. cholerae*; **Fig. 3B**; **Table S9**). Three of the top ten targets had chimeras with the *V. cholerae* homolog Vcr17 in Hfq RIL-seq data ^46^, indicative of direct base-pairing: VP0755, VP2480, and VPA0419. VPA0419 encodes a YfcZ/YiiS family protein of unclear function, while VP2480 and VP0755 encode proteins in the chitin utilization pathway (chitosugar ABC transporter component and chitobiase, respectively) ^53^ (**Fig. 3D**). Predictions revealed potential stable interactions between VcrX and these two mRNAs from *V. parahaemolyticus* over their ribosome binding sites, consistent with translational repression (**Fig. 3E**). A*s* chitin is central to *Vibrio* biology as a colonization surface, signal, and carbon source ^54^, and VcrX transcription may be activated by CRP-TfoX, we focused on these two targets.

Post-transcriptional repression of the *V. cholerae* VP0755 homolog (VC2217) was previously reported ^46^, and we confirmed that *V. parahaemolyticus* VcrX represses VP0755 in the same heterologous system (**Fig. S7B**). Unlike for *V. parahaemolyticus*, where we predicted one binding site, *V. cholerae* VcrX was predicted to interact with two binding sites in the *V. cholerae* VC2217 5’UTR (BS1, as well as BS2 over the translation initiation region ^46^; **Fig. S7C**). This suggests that specifics of the regulation of the chitobiase mRNA may have diverged between the two species.

Regulation of the VP2480 ABC transporter homolog VC0619 was not previously tested in *V. cholerae*, and we could not detect VP2480 GFP reporters in *E. coli* or *V. parahaemolyticus* to confirm post-transcriptional control. The gene is encoded in a long operon with an unclear TSS based on our data (**Fig. S7D**). RNA-seq after VcrX pulse-overexpression in M9 + GlcNAc(2-6) to induce target expression showed mild (<2-fold) but significant repression of VP2480 and several downstream genes (**Fig. S7E; Table S10**). This level of differential expression was surprising, as highly stable base-pairing over the translation initiation site is conserved across the *Vibrio* genus and even into *Aliivibrio* (**Fig. S7F**). VP2480 regulation might thus be mostly visible at the translational level, or under specific conditions not tested here.

VcrX therefore represses at least two chitin-related genes, and this regulation appears to be conserved in *Vibrio*. To determine if repression might have phenotypic consequences, we tested if VcrX overexpression could affect growth of *V. parahaemolyticus* in M9 with GlcNAc(2-6) as the sole carbon source. To force utilization through the targets VP0755 and VP2480 (especially as VP0755 may not be essential for growth on chitin due to functional redundancy ^53^), strains were generated in a Δ*nagE* background (VP0831, **Fig. 3D**). The Δ*nagE* strain carrying the wild-type VcrX plasmid had a growth defect in GlcNAc(2-6) only when IPTG (isopropyl-β-D-1-thiogalactopyranoside) was added to induce sRNA expression, and this phenotype was rescued by the M1 mutation in the VP0755/VP2480 seed region of the sRNA (**Fig. 4F**). Therefore, VcrX regulation can affect *V. parahaemolyticus* use of chitosugars.

**Figure 4.**
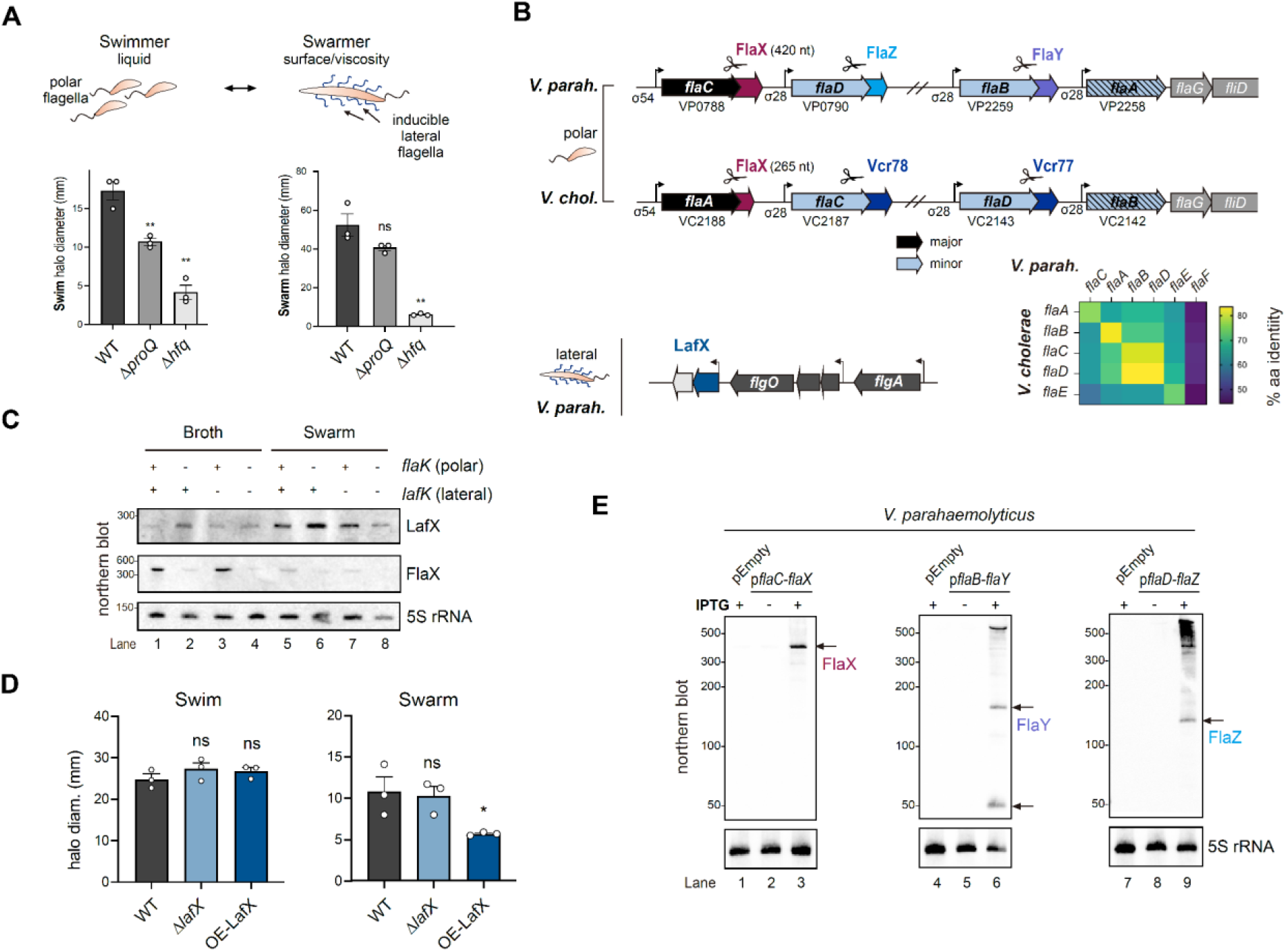
Conserved and specific *V. parahaemolyticus* flagella related sRNAs. **(A)** (*Top*) Two modes of *V. parahaemolyticus* motility driven by independent flagellar systems. (*Bottom*) Effect of Δ*hfq/*Δ*proQ* on *V. parahaemolyticus* polar and lateral motility phenotypes in RIMD 2210633: swimming and swarming. n=3 ** p<0.01. ns: not significant, mean vs. WT. **(B)** Context of sRNAs encoded at flagellar gene loci in *V. parahaemolyticus* and *V. cholerae. V. parahaemolyticus* candidate LafX and the adjacent lateral flagella cluster are absent in *V. cholerae*. Black arrows: major polar flagellins. Light blue arrows: minor polar flagellins. Scissors: RNase E processing. Bent arrows: TSS. *Bottom right*: Homology between *V. cholerae* and *V. parahaemolyticus* major/minor polar flagellins. Percent amino acid identity was determined by BLASTp. **(C)** Northern blot of LafX/FlaX expression in broth (swim) and surface (swarm) conditions. FlaK/LafK: polar/lateral σ54-dependent regulators. **(D)** Effect of *lafX* deletion/overexpression on RIMD 2210633 swimming/swarming. *p<0.05. ns: not significant, vs WT. Student’s *t*-test, n=3. **(E)** Northern blot of *V. parahaemolyticus* 3’ UTR-derived flagellar sRNAs expressed with parental flagellin transcript from an IPTG-inducible promoter. For northern blots, 5S rRNA served as a loading control.

### Conserved and species-specific flagella-associated sRNAs

Flagella are major drivers of environmental adaptation, affecting motility, host/surface interactions, and phage susceptibility. We wondered if sRNAs integrated into the *Vibrio* flagellar systems might provide an opportunity to study the dynamics of sRNA networks during adaptation. *V. parahaemolyticus* is a model for studying motility, including the hierarchical transcriptional cascades that regulate flagellar biosynthesis, and encodes two independent flagellar systems (polar and lateral; **Fig. 4A**) - unlike *V. cholerae* - providing an additional opportunity to identify species-specific sRNAs. Absence of Hfq and ProQ affected both swimming (polar flagella) and swarming (lateral flagella) of *V. parahaemolyticus* strain RIMD 2210633, suggesting sRNAs might regulate both systems.

To identify such regulators, we searched for those encoded near motility genes, which identified four candidates (**Fig. 4B**): three sRNAs encoded downstream of polar flagellins (VPnc0022/FlaX, VPnc0023/FlaZ, VPnc0047/FlaY; **Fig. S8A**), and VPnc0080 (“LafX”; **Fig. S8B**), which is encoded in a lateral flagella gene cluster. The lateral system encodes environmentally regulated, inducible peritrichous flagella, which mediate surface swarming in RIMD 2210633 (**Fig. 4A**). LafX is conserved in Harveyi clade species that encode lateral flagella (**Fig. 2**; **Fig. S8C**), and absent in *V. cholerae* and species that lack this secondary system. While BLAST identified a potential match in *V. natriegens* (**Fig. 2**), which lacks lateral flagella, it was unrelated to the region downstream of *flgO* that encodes LafX in the other species. LafX was more highly expressed in bacteria harvested from swarm plates than from broth but was not dependent on the lateral σ54-dependent regulator LafK (**Fig. 4C**). While LafX overexpression reduced swarming, but not swimming, motility (**Fig. 4D**), target predictions (IntaRNA ^52^) did not reveal flagellar mRNA targets. Thus, how LafX affects swarming is currently unclear.

The polar flagellar filament expressed by all Vibrio comprises a single σ54-dependent major flagellin, as well as a variable number of downstream σ28-dependent minor flagellins (*e.g*., five in *V. cholerae*, six in *V. parahaemolyticus*) ^56–58^. These flagellins, including the co-induced σ28-dependent minor flagellins, are differentially inserted along the filament, hinting at post-transcriptional control. We identified an abundant sRNA, VPnc0022, encoded in the 3’ UTR of the major polar flagellin gene *flaC* (**Fig. 4B**). We subsequently found that VPnc0022 shares homology and genomic context with *V. cholerae* FlaX, which is processed from the *flaA* major flagellin mRNA by RNase E ^35,59^ (see **Fig. 4B**, *bottom inset* for flagellin homology/nomenclature between the two species). However, *V. parahaemolyticus* FlaX has an ∼160 nt extension at its 5’ end compared to other *Vibrio* species, which was not previously reported (**Fig. S9A, B**) ^35^, and we confirmed the expression of a longer FlaX transcript of >300 nt (**Fig. 4C**; **Fig. S6B**). A putative sORF of 12 codons is encoded in the inserted region (**Fig. S9B**), but we so far have not detected its translation.

We also identified two candidate sRNAs encoded in the 3’ UTRs of minor flagellin mRNAs (**Fig. 5B; Fig. S8A**): VPnc0023 (*flaD*/VP0790 3’ UTR) and VPnc0047 (*flaB*/VP2259 3’ UTR). cDNA library coverage suggested that both, like FlaX, are processed from their parental transcripts (**Fig. S8A**). Two sRNAs derived from minor polar flagellin 3’ UTRs (Vcr77 and Vcr78) have also been reported in *V. cholerae* ^35^. VPnc0023 shows weak homology to Vcr77, although most homology arises from regions overlapping with the flagellin ORF (**Fig. S10A**). VPnc0047 has limited homology to the Vcr78 terminator. Neither sRNA shows similarity to FlaX, as was proposed for Vcr77/78 ^35^. We renamed VPnc0047 and VPnc0023 as FlaY and FlaZ, respectively, in the order we identified them.

**Figure 5.**
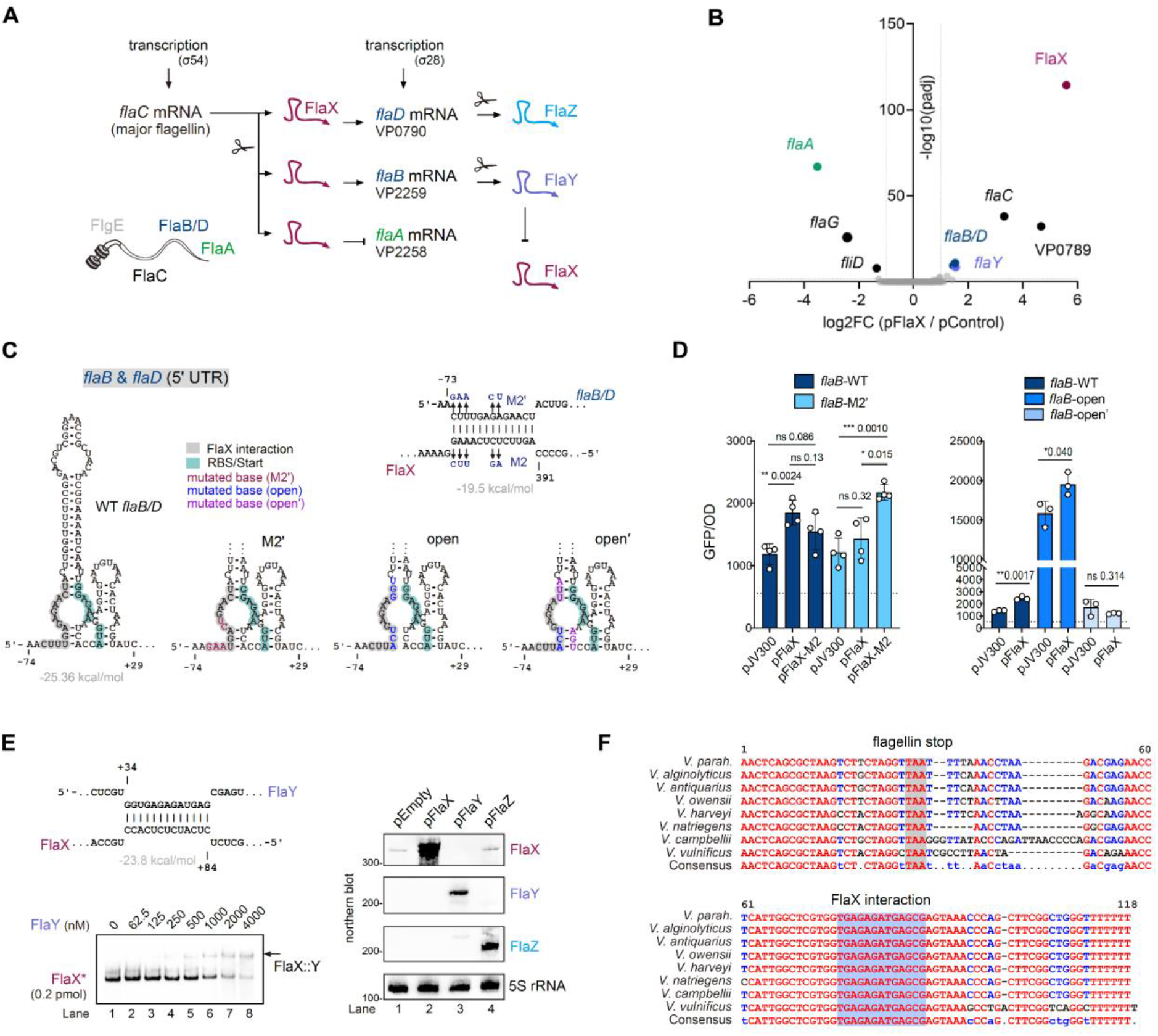
Post-transcriptional regulation of *V. parahaemolyticus* polar flagellins. **(A)** Working model for *V. parahaemolyticus* polar flagellin regulation based on this and previous ^35^ work in *V. cholerae*. **(B)** RNA-seq differential expression after FlaX pulse-overexpression vs. empty vector control in MLB. n=2 independent cultures. Dashed lines: log2FC = |1|; -log10(p) = 2. **(C)** (*Top right*) Predicted interaction (IntaRNA ^52^) between homologous *flaB* (VP2259) / *flaD* (VP0790) 5’ UTRs and FlaX. (*Bottom*) predicted *flaB/D* 5’ UTR secondary structure (RNA-fold ^60^). **(D)** GFP translational reporter levels for *E. coli* two-plasmid system analysis of *flaB* translation in the absence/presence of FlaX WT and point mutants in the interaction (*left*) or secondary structure (*right*) regions. *p<0.05, **p<0.01, ***p<0.001, ns: not significant, vs WT. Student’s *t*-test, n=3. **(E)** (*Top left*) Predicted base-pairing between FlaY and FlaX. (*Bottom left*) Electrophoretic mobility shift assay with *in vitro*-transcribed FlaX (biotin labeled) and FlaY. Representative of three independent experiments. (*Right*) Northern blot of FlaX after pulse-overexpression of FlaY or FlaZ. **(F)** Alignment of VP2259/*flaB* homolog 3’ UTRs (*flaY*). BLASTn matches from *Vibrio* species with coverage >90% and percent identity >80% were included. Sequences were aligned with multalin ^61^. See **Fig. S10B** for extended alignment.

Consistent with cotranscription and processing from σ54-dependent *flaC*, FlaX levels were dependent on the polar σ54-dependent activator FlaK (homolog of *V. cholerae* FlrA) (**Fig. 4C**). Expression of FlaY and FlaY was below the limit of northern blot detection even in WT under the conditions tested. To confirm that the three sRNAs are processed from their parental mRNAs, we cloned the flagellin genes *flaC*, *flaB*, and *flaD* with their respective 3’ UTRs behind an IPTG-inducible promoter on a plasmid. Consistent with co-transcription and processing, all three sRNAs were only detected in the presence of the inducer (**Fig. 4E**). FlaX expressed from this context was similar in length (>300 nt) as what we detected from the native context in WT. From this context, we detected two FlaY species (∼150 nt/50 nt) and a single version of FlaZ (<150 nt). While homology is limited between FlaY/Vcr78 and FlaZ/Vcr77, northern blot patterns between *V. parahaemolyticus* and *V. cholerae* were similar, with two bands visible for Vcr78/FlaY (^29^, **Fig. 4E**).

While these observations suggest that *V. parahaemolyticus*, like *V. cholerae*, encodes three sRNAs processed from hierarchically-transcribed flagellin mRNAs, we observed substantial diversity in sequence and length, especially for those derived from minor flagellin sRNAs. Therefore, it is unclear if the sRNAs are evolutionarily related or if they serve a similar function in each species.

### Differential regulation of polar flagellins by FlaX and species-specific post-transcriptional feedback

In *V. cholerae,* σ54-regulated FlaX represses translation of the σ28-regulated minor flagellin mRNA *flaB* (VC2142, homolog of *V. parahaemolyticus flaA*/VP2258; **Fig. 4B**) ^35^. This suggests that the sRNA is integrated into the hierarchical transcriptional cascade of polar flagella (**Fig. 5A**). However, how FlaX, as well as the two minor flagellin sRNAs, coordinate the *V. cholerae* polar cascade was not fully clear. To identify regulatory targets of *V. parahaemolyticus* FlaX, we pulse-overexpressed the sRNA and measured differential expression compared to an empty-vector control strain by RNA-seq. Strikingly, all differentially expressed transcripts encoded polar flagella genes. While *flaA-flaG-fliD* (VP2258-VP2256) was lower after FlaX induction, *flaB* and *flaD* (VP2259 and VP0790) mRNAs were increased (**Fig. 5B; Table S11**). In line with VP2259 upregulation, levels of *flaY* were also increased 2-fold.

The flagellin encoded by *flaA,* which was repressed by *V. parahaemolyticus* FlaX, is most closely related in sequence and genomic context to *V. cholerae flaB* (**Fig. 4B**, *inset*), which is also repressed by FlaX ^35^. This suggested that this aspect of FlaX regulation might be conserved between the species. Consistent with this, we predicted three stable binding sites (BS) between *V. parahaemolyticus flaA* and FlaX, including one over the ribosome binding site, as was confirmed for *V. cholerae* homolog *flaB* (**Fig. S11A**) ^35^. Using a two-plasmid reporter system ^62^, we confirmed that *V. parahaemolyticus* FlaX represses *flaA* post-transcriptionally, while having no effect on translation of upstream *flaC* (**Fig. S11B**). The FlaX and *flaA* 5’ UTR regions involved in these interactions are also conserved in *Vibrio* (**Fig. S9B, S11C**), as previously reported ^35^. Therefore, repression of this flagellin by FlaX appears to be a shared mechanism.

Pulse overexpression suggested that FlaX might also activate the minor flagellin genes *flaB* and *flaD* (**Fig. 5B**). In line with this, the second most abundant ProQ RIL-seq interaction partner for *V. cholerae* FlaX was *flaD*/VC2143 ^35^ (corresponding to *flaB*/VP2259 in *V. parahaemolyticus*; **Fig. 4B**, *inset*). While reporter assays suggested that *V. cholerae* FlaX activates *flaD* translation, the mechanism was not explored. We predicted an interaction between *V. parahaemolyticus* FlaX and the identical *flaB* / *flaD* 5’ UTRs, as well as a potential secondary structure involving the FlaX targeting site that could hinder translation initiation (**Fig. 5C**), as reported for other targets activated by sRNAs ^63^. Using the *E. coli* two-plasmid system, we confirmed that FlaX post-transcriptionally activates the *flaB*/*D* 5’ UTR via base-pairing (**Fig. 5D**, *left*). Moreover, while mutations in the predicted secondary structure (**Fig. 5C**) increased reporter expression, compensatory mutations that would restore base-pairing returned GFP to WT levels, consistent with a structure inhibiting translation (open/open’; **Fig. 5D**, *right*). The mutations, which are outside the FlaX pairing region (**Fig. 5C**), also reduced upregulation by the sRNA. Although the mechanism requires further examination, this structure is conserved, suggesting it has an important function (**Fig. S11D, E**).

Based on conservation, predictions, and experimental data in *V. cholerae* and *V. parahaemolyticus* taken together, we expand on the current model of FlaX regulation (**Fig. 5A**). σ54-dependent FlaX is co-transcribed with the major flagellin FlaC, processed, and differentially up/downregulates σ28-dependent minor flagellins to fine-tune their induction. This differential regulation is nicely consistent with the model of spatial flagellin distribution in *V. cholerae* based on fluorescent tagging, where the σ54-dependent major flagellin is incorporated immediately after the hook, followed by differential incorporation of the co-transcribed σ28-dependent minor flagellins: FlaX-activated FlaC/D, followed by FlaX-repressed FlaB towards the tip ^56^. Also supporting this model, *V. cholerae flaX* mutant strain filaments have different flagellin compositions compared to WT, as well as a swimming phenotype ^35^.

We next wondered about the role of the two σ28-dependent sRNAs (FlaY and FlaZ in *V. parahaemolyticus*) in this model. Recovery of Vcr77 and Vcr78 chimeras with minor flagellin mRNAs in *V. cholerae* ProQ RIL-seq suggested they could also directly activate or repress minor flagellin gene expression in *V. cholerae* ^35^. However, these sRNAs are not well conserved between *V. cholerae* and *V. parahaemolyticus*, and IntaRNA also did not identify flagellin mRNAs as potential targets for either FlaY or FlaZ in *V. parahaemolyticus.* Strikingly, however, we noticed that FlaY might base-pair with FlaX (**Fig. 5E**, *top left*). Interestingly, a similar interaction was not predicted between *V. cholerae* FlaX and Vcr77 or Vcr78 using the same parameters but FlaY - and particularly the region predicted to base-pair with FlaX - is well conserved in many other *Vibrio* species (**Fig. 5F; Fig. S10B**). Electrophoretic mobility shift assays showed that FlaX and FlaY can interact *in vitro* (**Fig. 5E**, *bottom left*).

We hypothesized that FlaY could provide a layer post-transcriptional feedback control in the system by sequestering FlaX activity, or alternatively by influencing its expression. To test this, we overexpressed FlaY or FlaZ from an IPTG-inducible promoter and measured levels of native FlaX by northern blot (**Fig. 5E**, *right*). FlaX was undetectable when FlaY, but not FlaZ, was overexpressed. This suggests that FlaY can affect FlaX expression or stability, and that some *Vibrio* species may have evolved post-transcriptional feedback in the polar cascade (**Fig. 5A**).

### *V. parahaemolyticus* encodes conserved and species-specific dual function sRNAs

Bacterial sRNAs can be “dual-function”, acting both as base-pairing regulators and as template for the translation of small proteins that work in related pathways ^2^. So far in *Vibrio*, one example, VcdR (**Fig. S12A**), has been described. In *V. cholerae*, VcdR base-pairs with several transcripts related to metabolism and encodes a small protein, VcdP (29 aa) that binds and regulates citrate synthase ^40^. In *E. coli,* Spot 42 also encodes a small protein, SpfP, which binds and regulates CRP, serving as a dual-function regulator of carbon metabolism and uptake genes ^64^. *Vibrio* Spot 42 also encodes a putative sORF with 12/15 amino acid residues, including key His10, shared with *E. coli* (**Fig. 6A, B**; **Fig. S12B**). The *Vibrio spfP* translation initiation site is likewise tied up in a secondary structure (**Fig. S12C**). We confirmed that *V. parahaemolyticus vcdP* and *spfP* can be translated with a GFP fusion (**Fig. 6C**). Thus, the dual-function property of Spot 42 is conserved outside the Enterobacteriaceae.

**Figure 6.**
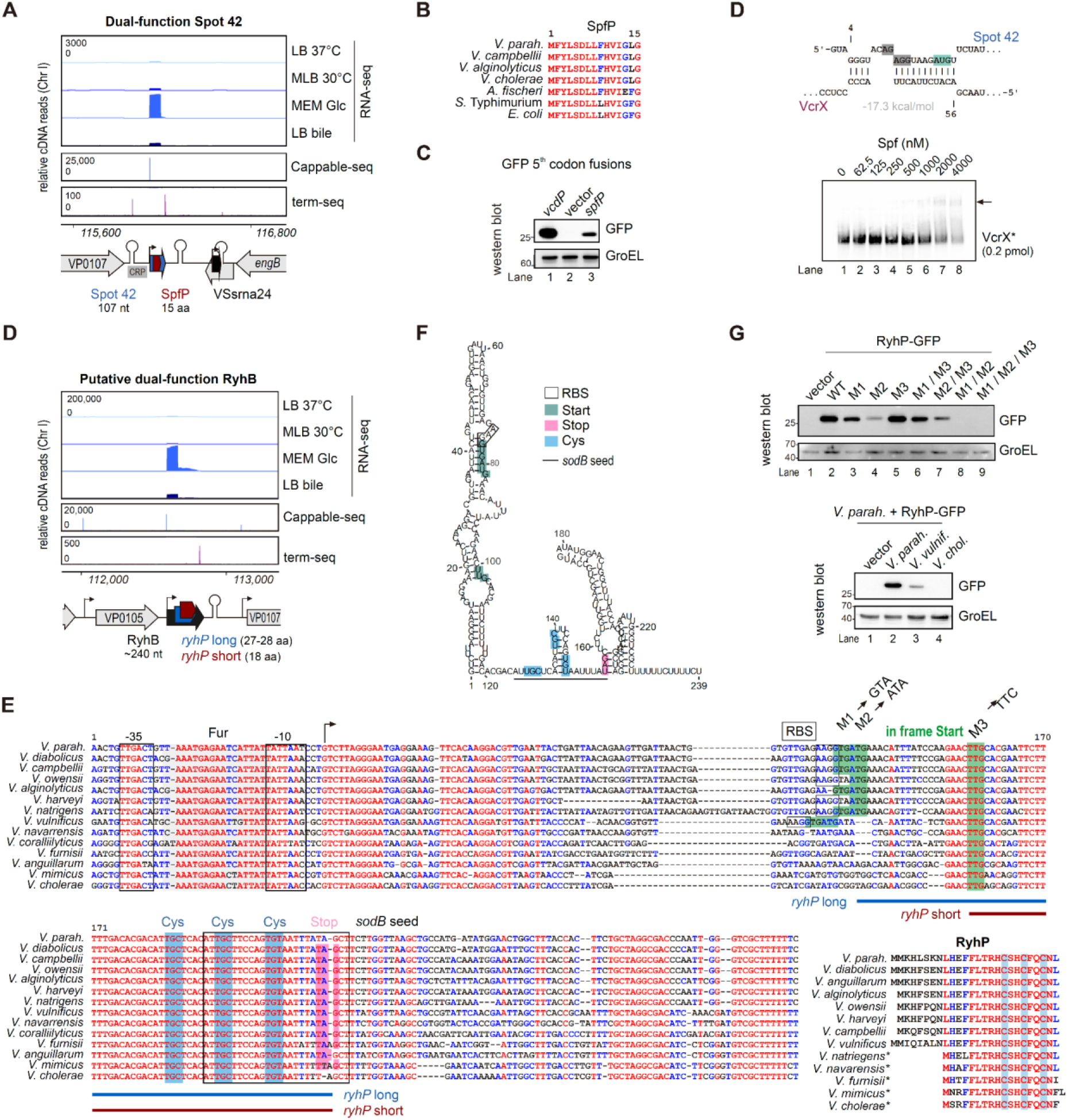
Dual-function Spot 42 and RyhB. **(A)** RNA-seq coverage for *V. parahaemolyticus* Spot 42. Blue: sRNA. Red: sORF. **(B)** Protein sequence alignment of SpfP homologs from *Vibrio* and Enterobacteriaceae species. **(C)** Western blot of *spfP* and *vcdP* translational GFP fusions. GroEL: loading control. **(D)** (*Top*) Predicted interaction between Spot 42/VcrX. Grey: ribosome binding site. Green: start codon. (*Bottom*) Gel mobility shift assay between Spot 42 and biotin-labeled VcrX (*). **(E)** RNA-seq coverage for potential dual-function RyhB. Black: sRNA. Blue/red: long/short *ryhP*. **(E)** Alignment of *ryhB* regions from *Vibrio* species. M1-M3: Point mutations in potential start codons tested in *V. parahaemolyticus*. (*Inset, bottom right*) RyhP protein sequence alignment for homologs identified in *Vibrio* species. *: may not be translated. Alignments were generated with multalin ^61^. **(F)** Predicted secondary structure of *V. parahaemolyticus* RyhB using RNAFold ^60^. **(G)** Western blots of *ryhP* GFP translational fusion expression in *V. parahaemolyticus*.

Above, we noted an inverse relationship between Spot 42 and VcrX levels, which could be explained by differential transcriptional control by CRP (**Fig. 3C**). However, we predicted base-pairing between the two sRNAs and observed binding between them *in vitro* (**Fig. 6D**). While the two sRNAs might affect each other’s stability and/or activity, as reported for other “sponge” sRNAs, an alternative hypothesis is that VcrX regulates translation of the SpfP small protein, as VcrX base-pairing over the *spfP* translation initiation region is conserved in *V. parahaemolyticus* and *V. cholerae* (**Fig. 6D**; **Fig. S12D**).

We next wondered if we could identify dual-function RNAs that have not yet been reported. One candidate that caught our attention is the *Vibrio* homolog of the iron-related sRNA RyhB, which is functionally analogous to its *E. coli* counterpart but much longer (∼240 vs. 90 nt) ^65–67^. It has also been suggested that *E. coli* RyhB might be translated, but this has not been validated ^68^. We identified three potential sORFs on *V. parahaemolyticus* RyhB that share the same stop codon (28, 27, 18 amino acids) (**Fig. 6D, E**). The 3’ end of all three overlaps the *sodB* pairing region (**Fig. S12E**), and as for *spfP*, the RBS of the longer ORF as well as start codons for all three are potentially located in a base-paired stem (**Fig. 6F**).

To confirm that *Vibrio* RyhB is translated, we fused its TSS up to the first five codons (with respect to the shortest sORF) to GFP on a plasmid. Western blot of *V. parahaemolyticus* carrying a 5th codon GFP fusion confirmed translation (**Fig. 6G**, *top*). As there are three potential start codons in-frame with the shared stop codon, we introduced point mutations into each (**Fig. 6E**). This revealed that the second ATG start codon likely drives most translation, with the first GTG codon also contributing (possibly as part of the RBS) (**Fig. 6G**, *top*). Interestingly, the more widely conserved TTG start codon was not used. In addition, *V. cholerae* RyhB does not encode the first two start codons or the RBS (**Fig. 6E**), and a *V. cholerae* GFP fusion was not translated, in contrast to the *V. parahaemolyticus* and *V. vulnificus* versions (**Fig. 6G**, *bottom*). Together, this suggests that RyhB is a dual function sRNA, but that this feature is restricted to only some *Vibrio* species. We term the sORF *ryhP* following previous nomenclature ^40,64,68^, although it does not resemble the predicted *E. coli* sORF.

*Vibrio* RyhB, like the Enterobacteriaceae homolog, is induced in response to iron limitation and regulates iron homeostasis and oxidative stress transcripts ^65,66,69^. The dual-function property of RyhB may have arisen within specific *Vibrio* lineages to, *e.g.* fine-tune the iron-sparing response or even interconnected pathways such as iron-sulfur cluster homeostasis. Interestingly, RyhP conserves three Cys residues in a HCxHCxxC motif, which could facilitate metal ion (Fe, Zn) or Fe-S interactions (**Fig. 6E**, *inset bottom right*). The protein is also His-rich with a basic pI of 8.7.

## Discussion

Here, we generated a high-resolution transcriptome and curated sRNA resource for *Vibrio parahaemolyticus*, which provides a foundation for comparative analysis of sRNA-based regulation across *Vibrio*. By combining transcriptome mapping, conservation analysis, and functional analysis, we identify conserved and taxon-specific sRNAs, demonstrate examples that regulate important genes and phenotypes, and uncover previously undescribed regulatory architectures. These include a dual-function RyhB, potential interplay between base-pairing and dual-function sRNAs, and sRNA–sRNA feedback restricted to a subset of *Vibrio* species.

This work provides not only a resource for *V. parahaemolyticus* and other *Vibrio* species, but also insight into how sRNA annotations can be transferred across a bacterial genus. We identified numerous shared sRNAs between relatively distant *V. parahaemolyticus* and *V. cholerae* (**Fig. 2**), particularly among characterized sRNAs, but overall overlap was limited. This observation is consistent with previous analysis in *V. splendidus*, where more than 60% of sRNAs were specific to the species ^45^.

Several factors may underlie the limited overlap. Some sRNAs are encoded on species-specific genomic islands or are associated with biological functions present in only some *Vibrio*: for example, TarAB is encoded on the *V. cholerae* pathogenicity island, while LafX is encoded with lateral flagella genes, which are absent from many *Vibrio*. Others may have diversified to suit the needs of regulatory networks remodeled to fit a new niche, analogous to independently-transcribed UhpU in *E. coli* ^70^. However, some apparent lineage-specific VPncRNAs may instead be spurious transcripts not under selection. Conversely, some sRNAs that are in fact shared amongst more species may have escaped detection because of low expression in examined conditions, or rapid sequence divergence.

Convergent evolution can also generate functional homologs that share targets but not sequence homology. Comparative experimental approaches such as RIL-seq, which couples sRNA discovery to target identification, applied to different strains or species would identify functional homologs through shared target spectra rather than sequence similarity. This might uncover functional or highly diverged homologs that we missed, including a counterpart of *V. cholerae* QrrX ^46^. Moreover, as an sRNA with the same sequence can in principle have different expression patterns or target spectrum, these experimental approaches, coupled with lab validation, are crucial for accurately describing the divergence of sRNAs between groups.

Notably, several *Vibrio* 3′ UTR-derived sRNAs showed divergences between species, such as VPnc0074 (absent in *V. cholerae*), OueS (absent in *V. parahaemolyticus*), and FlaY/Z (divergent in sequence and potentially function between the species). While additional experimental data are needed to confirm these observations, they are in line with the proposal that 3′ UTRs are substrates for sRNA evolution ^5,44^. Moreover, *Vibrio,* and for example divergent FlaY and FlaZ, might serve as a framework for studying this process at different taxonomic scales ^71^.

The most striking sRNA in our annotation was highly conserved and abundant VcrX. VcrX expression was inversely correlated with Spot 42 across several conditions and was dependent on CRP. Although the conserved motif in the VcrX promoter is not a perfect CRP-S site, it might also require the chitin-responsive co-activator TfoX ^51^. This is consistent with conserved direct regulation of two chitin-related transcripts across *Vibrio*. However, VcrX repression of chitin utilization genes raises the question of why the sRNA might be induced by chitin via TfoX. One possibility is that VcrX separates the induction of chitin utilization genes from competence genes that are co-regulated transcriptionally by TfoX. Alternatively, if targets VP0755 and VP2480 are activated by the same transcription factor as VcrX, the sRNA could form part of an incoherent feed-forward loop that modulates dynamics of chitin gene induction. However, as VcrX may have additional mRNA targets ^46^, a dedicated role in repressing chitin mRNAs might be an oversimplification.

Also complicating matters is the potential for VcrX to regulate Spot 42, and potentially *spfP* translation. This would potentially impact CRP and its regulon, if the function of SpfP is shared with *E. coli* ^64^, as well as targets of Spot 42 base-pairing. Thus, the interplay between these sRNAs may be more complex than differential CRP repression/activation. While RNA-sRNA antagonism has recently been described in *Vibrio* ^46^, regulation of dual-function sRNA stability or translation would be a new addition to the bacterial post-transcriptional regulation toolkit. Overall, VcrX may be part of a complex regulatory network dedicated to *Vibrio* carbon source utilization, and could drive further study of metabolic and RNA network architecture ^72^.

As chitin is central to *Vibrio* biology as an important carbon source and environmental signal ^50^, it is not surprising that multiple sRNAs that respond to chitin or regulate chitin genes have been identified: TfoR, MicX, VcdR, and now VcrX ^32,40,73^. Interestingly, *Vibrio* appears to lack a homolog of ChiX, which in *Salmonella* is a repressor of *chiPQ* mRNA and is antagonized by an intergenic fragment of *chb* mRNA ^74^. Whether *Vibrio* encodes a similar “decoy” scheme awaits full characterization of these sRNAs.

Although hierarchical transcriptional control of flagellar biosynthesis has long been a model of bacterial gene regulation, the contribution of sRNAs to this cascade has only recently become apparent ^35,75,76^. We found that as in *V. cholerae* ^35,59^, *V. parahaemolyticus* encodes three sRNAs derived from polar flagellin mRNA 3′ UTRs: a longer homolog of FlaX (VPnc0022), and two (FlaY/VPnc0047, FlaZ/VPnc0023) that are encoded in the same context as Vcr77 and Vcr78, but do not share homology outside of regions overlapping with flagellin ORFs. We also identified an sRNA encoded adjacent to lateral flagella genes, VPnc0080 (LafX), that influences swarming by an unknown mechanism. LafX carries a differentiated single-nucleotide polymorphism (SNP) in the *V. parahaemolyticus* Molassodon ecospecies, which has distinct lateral flagella that are regulated differently from Typical strains ^22^.

For FlaX, we confirmed that previous findings in *V. cholerae* apply across the genus and also expanded on and clarified its role in regulating polar flagellins ^35,59^. Processing from the σ54-dependent major flagellin mRNA and repression of a σ28-dependent minor flagellin (*flaA* in *V. parahaemolyticus*) is likely conserved in many *Vibrio* species. We also provide evidence that FlaX directly activates two additional minor flagellins by disrupting a conserved structure occluding their RBS, both in *V. parahaemolyticus* and *V. cholerae*. Together, these findings support a model (**Fig. 5A**) in which FlaX fine-tunes temporal induction of three minor flagellins, which are co-induced transcriptionally by σ28, to facilitate differential incorporation into the filament ^56^. This is supported by different filament compositions in *V. cholerae* FlaX strains and is an elegant mechanism to temporally and spatially control expression and incorporation of the different flagellins into the filament.

While FlaX regulation of minor flagellins appears conserved in *Vibrio*, elements downstream may have diversified to accommodate differences in flagellar structure and function. As in *V. cholerae*, we identified two sRNAs processed from σ28-dependent minor flagellin 3′ UTRs, but these are not obviously homologous. Notably, FlaY in *V. parahaemolyticus* and several other non-*cholerae* species conserves a potential FlaX-pairing region. We hypothesize that FlaY may be a FlaX sponge, adding an additional layer of feedback to the cascade. The role of FlaZ remains unclear.

Bacterial genomes likely encode unannotated small proteins ≤50 amino acids ^77^, including on so-called “dual-function” sRNAs. Here, we demonstrate that 15 aa SpfP, first identified on *E. coli* Spot 42 ^64^, is also translated in *Vibrio*, providing taxonomic evidence that SpfP has an important cellular role. Moreover, the *Vibrio* small protein conserves key residues involved in CRP interaction in *E. coli*, suggesting this role is also conserved. Only ∼10 dual-function sRNAs are so far known, and we found that RyhB from some *Vibrio* species can be added to the list. Our discovery of *ryhP* highlights the value of studying sRNAs in different species.

A function for RyhP remains to be identified. The sORF is enriched in Cys codons, which fits with the role of RyhB in iron- and Fe-S cluster homeostasis ^65,67^. The three Cys residues are part of a HCxHCxxC sequence, which could mediate metal binding or be redox-sensitive. The small protein could bind an iron transporter, or potentially a regulator like Fur, reinforcing, antagonizing, or subdividing its regulation of target promoters. Translation of the sORF could also be responsive to cellular Cys levels. The *ryhP* coding sequence overlaps the conserved *sodB* seed region, as for *E. coli* Spot 42 (and AzuR) ^64,78^. The translation initiation region of *ryhP*, also like Spot 42, appears to be in a structured region. Thus, RyhB might provide an avenue to explore the interplay between the mRNA and sRNA roles of dual-function RNAs. Moreover, it suggests that even more examples of dual-function sRNAs might be hiding in plain sight.

## Materials and methods

### Bacterial strains and general culture conditions

*E. coli* and *Vibrio* strains used in this study are listed in **Table S12**. Recombinant strains were generated using plasmids and oligonucleotides listed in **Tables S13** and **S14** as described below and in the **Supplementary Methods**. Key sequences (cloned regions, point mutations) are listed in **Table S15**. For cloning, *E. coli* DH5α or DH5α λ*pir* was used. *E. coli* SM10 λ*pir* or MFD*pir* were used for *Vibrio* plasmid conjugation. For RNA-seq transcriptome mapping, the available environmental *V. parahaemolyticus* VN-5003-Wue strain, a version of VN-5003 (SAMN03470171) obtained from Prof. Kai Papenfort was used. All additional lab work and RNA-seq was performed with *V. parahaemolyticus* RIMD 2210633 ^28^. *E. coli* strains were routinely grown in LB broth at 37°C with shaking or LB plates with 1.5% agar, with addition of selective antibiotics as required at the following concentrations: carbenicillin, 100 μg/ml; kanamycin - 50 μg/ml; chloramphenicol - 25 μg/ml. *Vibrio* strains were routinely grown in LB or MLB (LB with 3% NaCl) broth with shaking or agar, at 37°C or 30°C, respectively, with appropriate antibiotics at the following concentrations: chloramphenicol - 5-10 μg/ml, kanamycin - 250 μg/ml; spectinomycin - 50 μg/ml.

### Growth conditions, RNA isolation, cDNA library preparation, and data analysis

Extended details for sample preparation and data analysis of RNA-seq analysis in strain VN-5003-Wue are provided in the **Supplementary Methods.** For generation of RNA-seq, dRNA-seq, Cappable-seq, and term-seq in the wild-type strain VN-5003-Wue, bacteria were grown overnight in MLB broth at 30°C. For the LB37 and MLB30 condition, cells were diluted 1/100 into fresh LB or LB with 3% NaCl and grown at 37°C or 30°C, respectively, until log phase (OD_600_ approx. 0.5) (LB37 and MLB30 conditions). T3SS1- and T3SS2-inducing conditions were generated as described previously ^30^. For the T3SS1 condition, cultures growing in LB at 37°C in log phase were washed 3 times in DMEM (Gibco, MEM + Glc), inoculated into fresh DMEM, and then incubated for 1 hour at 37°C. For the T3SS2 condition, a log phase culture in LB at 37°C was supplemented with 0.05% ox bile salts (Sigma) for 1 hour. Transcription was halted by the addition of 0.2 volumes of Stop mix (95% ethanol/5% phenol) and immediate freezing in liquid N_2_. Cells were stored at -80°C until processing. RNA was extracted using hot phenol and libraries were prepared by by Vertis biotechnologie (Freising, Germany) according to their protocols and sequenced on a NextSeq500 instrument (high-output, 75 cycles) at the Core Unit SysMed at the University of Würzburg.

### Small RNA pulse-overexpression and transcriptomics

To identify putative RNA targets of FlaX, we grew cultures of WT carrying pZJ14 (empty) or pSSvSH87 (FlaX under control of the pBAD promoter) until log phase in MLB at 30°C. Arabinose was added to cultures for 20 minutes at a final concentration of 0.1%. To identify putative targets of VcrX, we grew cultures of WT carrying either pMMB207 (empty plasmid) or pSSvSH131 (VcrX under control of the Ptac promoter in pMMB207) in M9 medium with 1% NaCl and 0.2% GlcNAc(2-6) as the sole carbon source at 30°C until log phase. IPTG was then added to a final concentration of 1 mM for 20 min. In all cases, cultures were mixed with Stop mix at the end of the experiment and immediately frozen at -80°C. RNA was extracted as described above, and subjected to DNase I digestion, library preparation, and sequencing at Novogene (Beijing). Data analysis was performed as described in the **Supplementary Methods.**

### sRNA conservation

Reference genomes of 22 species (**Table S16**) were downloaded from NCBI. BLASTn was used with the *V. parahaemolyticus* RIMD 2210633 homolog as query with the following parameters: task=“blastn”, word_size=4, qcov_hsp_perc=30, perc_identity=30, evalue=1000000, gapopen=0, gapextend=2, penalty=-1, reward=1, max_target_seqs=50000. We calculated a “Conservation Index” for each hit by taking the square root of the product of Percent Identity and Query Coverage per HSP (High-scoring Segment Pair). We then kept the hit with the highest Conservation Index for each species. There was no hit with Percent Identity lower than 50. To identify sRNA homologs shared between *V. parahaemolyticus* and *V. cholerae*, we performed BLAST searches using our list of sRNAs (**Table S4**) and a list of published *V. cholerae* sRNAs ^29,34^. We used options “-task blastn -word_size 4 -qcov_hsp_perc 30 -perc_identity 30 - evalue 1000 -gapopen 0 -gapextend 2 -penalty -1 -reward 1 -max_target_seqs 5000” for BLASTn searching. For each *V. parahaemolyticus* sRNA, we kept the *V. cholerae* hit with the highest bitscore. Matches with E-value <0.1, coverage and percent identity >50% were retained. We also used BLASTn to compare *V. cholerae* sRNAs against *V. parahaemolyticus* genome using options “- task blastn -word_size 4 -qcov_hsp_perc 50 -perc_identity 50 -evalue 1000 -gapopen 0 - gapextend 2 -penalty -1 -reward 1 -max_target_seqs 5000”. For each *V. cholerae* sRNA, we kept the *V. parahaemolyticus* genome hit with the highest bitscore. Matches with E-value <0.1, coverage and percent identity >50% were retained.

### General recombinant DNA techniques

All mutant strains and plasmids used and/or constructed in this study are listed in **Tables S12 & S13**, respectively. Oligonucleotide primers (Sangon) are listed in **Table S14**. Restriction enzymes, *Taq* polymerase for validation PCR, and T4 DNA ligase were purchased from Tiangen, ThermoFisher Scientific, or New England Biolabs. Seamless cloning mix was purchased from Sangon. For cloning purposes, Phanta high-fidelity DNA polymerase was used (Vazyme). DNA constructs and mutations were confirmed by Sanger sequencing (Sangon). We generated point mutations via inverse PCR with overlapping mutagenic primers, *Dpn*I digestion, and validation via Sanger sequencing. The generation of *V. parahaemolyticus* deletion and point mutant strains, sRNA overexpression plasmids, and GFP reporters is outlined in detail in the **Supplementary Methods**.

### RNA extraction and northern blotting

Total RNA was extracted from approximately 4 OD_600_ of bacteria using the hot phenol method as described previously ^76^. RNA (10 µg) was separated on a 6% PAA (polyacrylamide) 7M urea gel, and then electrotransferred onto nitrocellulose membrane (GE). RNA was crosslinked to membranes (120 mJ) and then pre-hybridized for one hour at 42°C in Hybridization cocktails (Sangon). A 5’-end biotin labeled DNA oligonucleotide probe (Sangon) complementary to the RNA of interest was added (10 μl of 10 μM) and hybridized overnight at 42°C. Membranes were then washed for 15 min each with SSC + 0.1% SDS (5x, 1x, 0.5x) at 42°C. Membranes were then washed for 10 minutes at 37°C in 1x PBS, pH 7.4 + 0.5% SDS), blocked for 30 min at 37°C in blocking reagent (0.1% Tropix I-Block (Invitrogen), 0.5% SDS, 1x PBS, pH 7.4). Streptavidin-horseradish peroxidase conjugate (500 ng, Abcam ab7403) was then added for 1 hour at 37°C. After washing three times in PBS-SDS at 37°C, membranes were detected after the addition of ECL reagent (Sangon). In some cases, probes were labeled radioactively and blots were performed as described previously ^76^ or were detected with a 5’-DIG labeled probe essentially as described above.

### Electrophoretic mobility shift assays

RNAs of interest were *in vitro*-transcribed from templates with the T7 promoter sequence generated by PCR with Phanta Flash polymerase (VAzyme) using the TranscriptAid T7 High Yield Transcription Kit (Thermo Scientific) according to the manufacturer’s instructions. RNAs were then checked for quality by electrophoresis on a PAA-urea gel, Biotin labeling was performed by including dUTP-biotin (Beyotime) in reactions, followed by gel extraction. Gel shift assays were performed as described previously ^76^. Briefly, biotin-labeled RNA (0.04 pmol) was denatured (1 min, 95°C) and cooled for 5 min on ice. Yeast tRNA (1 μg, Invitrogen) and 1 μl of 10× RNA structure buffer (final concentration 10 mM Tris, pH 7, 100 mM KCl, 10 mM MgCl_2_) was then mixed with the labeled RNA. Unlabeled RNA (2 μl diluted in 1× structure buffer) was added to the desired final concentrations (0, 0.0625, 0.125, 0.25, 0.5, 1, 2, or 4 μM). Binding reactions were incubated at 37°C for 15 min. Before loading on a pre-cooled native 6% PAA, 0.5× TBE gel, samples were mixed with 3 μl native loading buffer (50% (v/v) glycerol, 0.5× TBE, 0.2% (w/v) bromophenol blue). Gels were run in 0.5× TBE buffer at 300 V and 4°C. Gels were blotted onto nitrocellulose membranes. After crosslinking (120 mJ), membranes were blocked (1× PBS, 0.5% SDS, 0.1% Tropix blocking reagent) for 30 minutes at 37°C. Streptavidin-horseradish peroxidase conjugate (AbCam) was added, and after 1 hour of incubation at 37°C, membranes were washed 3 times with 1x PBS, 0.1% SDS. Labeled RNA was visualized with enhanced chemiluminescence (ECL) reagent (Sangon). *In vitro*-transcribed template sequences are listed in **Table S15**.

### Protein analysis

GFP reporter translation was analyzed by fluorescence (pXG10 system) or SDS-PAGE and western blotting of whole-cell lysates (sORF translation). For western blot, the RIMD 2210633 WT strain carrying sORF-GFP fusion plasmids were grown in MLB with 5 ug/ml Cm at 30°C until mid-log phase (OD_600_ 0.4–0.5). IPTG was then added to a final concentration of 1 mM and cultures were incubated for an additional 30 minutes. Cells were harvested by centrifugation at 13,000 rpm. Cell pellets were resuspended in 100 μl of 1× protein loading buffer (62.5 mM Tris-HCl, pH 6.8, 100 mM dithiothreitol, 10% (v/v) glycerol, 2% (w/v) SDS, 0.01% (w/v) bromophenol blue) and boiled for 8 min. For analysis of total proteins, 0.05 OD_600_ of cells were loaded per lane on 12% SDS-polyacrylamide gels. Proteins were electro-transferred from gels to a PVDF (polyvinylidene fluoride) membrane. Membranes were blocked for 1 hr with 10% (w/v) milk powder in TBS-T (Tris-buffered saline-Tween-20) and then incubated overnight with primary antibody (anti-GFP, 1:1000, Roche #11814460001 in blocking reagent with BSA (Sangon; E661003-0200) at 4°C. Membranes were then washed with TBS-T, followed by 1 hr incubation with secondary antibody (anti-mouse IgG, Sangon D110087, 1:10,000 in BSA blocking reagent). After washing, the blot was developed using enhanced chemiluminescence reagent (Sangon) and imaged. As a loading control, a monoclonal antibody specific for GroEL (1:10,000; Sigma-Aldrich, #G6532-5ML) with an anti-rabbit IgG (1:10,000; Sangon, D110058) secondary antibody was used.

### Growth experiments

To test the effect of VcrX overexpression on growth with GlcNAc(2-6) as the sole carbon source, strains were first grown in pre-culture in 2 ml MLB at 30°C overnight with 5 μg/ml Cm. The next morning, cells were washed three times in PBS and inoculated into M9 with 1% NaCl and 0.2% GlcNAc(2-6) at an OD_600_ of 0.05 in duplicate. To one culture, IPTG was added to a final concentration of 1mM to induce VcrX expression. Cultures were grown at 30°C for 6 hours with shaking, and the final OD_600_ was read. The average of two independent experiments was plotted.

### Motility assays

Strains were grown in MLB until log phase, and then 0.05 OD_600_ was inoculated into swim or onto swarm plates. Swim plates (0.325% agar) and swarm plates (1.5% agar) were prepared as described previously ^30^. Plates were incubated at 30°C for approximately 16 hours until halos were observed. For diameter measurements, if the colony was not round, which means the longest diameter is more than 1 mm longer than the shortest diameter, we took the average between the longest and the shortest diameters.

### Data availability

All RNA-seq data have been deposited at GEO under accession GSE335512. The sequence of VN-5003-Wue (SAMN60828049) can be found at NCBI under BioProject PRJNA1478059. An updated annotation for RIMD 2210633 including sRNAs and TSS, as well as processed coverage files, can be found at https://doi.org/10.6084/m9.figshare.32682531.

## Acknowledgements

We thank Prof. Kai Papenfort for sharing the *V. parahaemolyticus* strain VN-5003-Wue, Qiyao Wang for sharing *V. parahaemolyticus* RIMD 2210633 and *E. coli* strains/plasmids, and Kim Orth for sharing plasmids. Laura Vogel, Huiqi Wen, and Philipp Kible provided technical assistance. We are grateful to Prof. Cynthia Sharma for support and valuable comments on the manuscript. We also thank members of the Chao and Sharma labs for fruitful discussions and feedback from Susan Gottesman on the manuscript. This research was supported by RFIS-II funding from the National Natural Science Foundation of China (NSFC, #32250610209) and funding from the German Research Foundation (DFG) under the framework of Priority Programme SPP2002 “Small Proteins in Prokaryotes: an Unexplored World”. Work in the DF laboratory is supported by an NSFC RFIS-III award (#32350710791) and the YC laboratory by the National Natural Science Foundation of China (#92478118, 32270064) and the Shanghai Municipal Science and Technology Commission (#24ZR1493200).

## Author contributions

Performed experiments: ZJ, HZ, SLS. Bioinformatic analysis: ZJ, HZ, SLS. Data analysis and interpretation: ZJ, HZ, SLS. Resources: DF, YC. Designed research: SLS. Writing: SLS. Editing: SLS, ZJ. Funding: SLS.

## Supplementary Methods

### Bacterial growth in four conditions for RNA-seq libraries

For generation of RNA-seq, dRNA-seq, Cappable-seq, and term-seq in the wild-type strain VN-5003-Wue, bacteria were grown overnight in MLB broth at 30°C. For the LB37 and MLB30 condition, cells were diluted 1/100 into fresh LB or LB with 3% NaCl and grown at 37°C or 30°C, respectively, until log phase (OD_600_ approx. 0.5) (LB37 and MLB30 conditions). T3SS1- and T3SS2-inducing conditions were generated as described previously ^16^. For the T3SS1 condition, cultures growing in LB at 37°C in log phase were washed 3 times in DMEM (Gibco, MEM + Glc), inoculated into fresh DMEM, and then incubated for 1 hour at 37°C. For the T3SS2 condition, a log phase culture in LB at 37°C was supplemented with 0.05% ox bile salts (Sigma) for 1 hour. Transcription was halted by the addition of 0.2 volumes of Stop mix (95% ethanol/5% phenol) and immediate freezing in liquid N_2_. Cells were stored at -80°C until processing.

### cDNA library generation, sequencing, and data analysis for normal RNA-seq

Total RNA was extracted from cell pellets using hot phenol as described previously ^17^. RNA was digested with DNase I and in some cases depleted of rRNA (Ribo-Cop Gram-negative, Lexogen). Libraries for RNA-seq, dRNA-seq, and term-seq were generated by Vertis biotechnologie (Freising, Germany) according to their protocols and sequenced on a NextSeq500 instrument (high-output, 75 cycles) at the Core Unit SysMed at the University of Würzburg. Alignment statistics for all cDNA libraries can be found in **Table S1.** Fastp ^18^ version 1.0.1 was used for quality control of the raw reads with default parameters. Reads were then mapped to the corresponding reference genome using Segemehl ^19^ version 0.3.4 with default parameters. FeatureCounts ^20^ version 2.1.1 was used for gene expression quantification with parameters “-f -s 1 -O --fraction --ignoreDup –primary”. Differential expression was calculated using R package DESeq2 ^21^ version 1.42.1 using the output from featureCounts without normalization. Other general sequencing file and alignment file manipulations (*e.g*., alignment statistics, genome coverage file generations, etc.) and calculations (*e.g*., TPM normalization) were performed using Samtools ^22^ version 1.22.1 and Julia ^23^.

### dRNA-seq transcription start site calling and sRNA predictions

Fastp ^18^ version 1.0.1 was used for quality control of the raw reads with default parameters. Reads were then mapped to the corresponding reference genome using Segemehl ^19^ version 0.3.4 with default parameters. Genome coverage files were generated using Samtools ^22^ version 1.22.1 and normalized using Julia ^23^. We employed the ANNOgesic pipeline ^24^ version 1.1.14 for TSS identification and sRNA prediction with TSS Predator 1.1beta. TSSs were identified using the “tss_ps” subcommand with default parameters. ANNOgesic also performed TSS classification, which followed the same rules as in ^25^. Primary TSSs were classified by having the highest coverage within 300 bp upstream (including the first base of the gene) of an ORF, and the other TSSs within this range were classified as secondary TSSs. Internal TSSs were classified as those located between the start and end of a gene (excluding the first base). Antisense TSSs were classified as locating on the opposite strand of the gene with 100 bp flanking regions. TSSs that did not belong to above classifications were identified as orphan TSSs. sRNA prediction was performed using the default parameters with options “--utr_derived_srna --filter_info tss blast_srna sec_str --compute_sec_structures” and with comparison to the Bacteria Small Regulatory RNA Database (downloaded from http://www.bac-srna.org/BSRD/index.jsp).

### Cappable-seq data processing and transcription start site calling

Fastp ^18^ version 1.0.1 was used for quality control of the raw reads with default parameters. Reads of the second replicate were reverse complemented. Both replicates were then mapped to the corresponding reference genome using Bowtie2 ^26^ version 2.5.5 with the “--local” option. We removed non-unique alignments by detecting “XS:i” in the optional field. Because reads were on the sense orientation, the leftmost position of the alignment matched the transcription termination site. Genome coverage using the leftmost base of each alignment was counted and normalized by counts-per-million using a customized Julia script using the XAM.jl package. Peaks were separately called for each replicate using Peaks.jl package with a width of 2. Peaks from the two replicates were merged with raw coverage values as peak heights. The heights of merged peaks were then normalized using the counts-per-million method. Rules of TSS classification were the same as for dRNA-seq, as defined in ^25^.

### Term-seq data analysis and termination site calling

Fastp ^18^ version 1.0.1 was used for quality control of the raw reads with default parameters. Reads were then mapped to the corresponding reference genome using Bowtie2 ^26^ version 2.5.5 with “--local” option. We removed non-unique alignments by detecting “XS:i” in the optional field. Because reads were on the antisense orientation, the rightmost position of the alignment matched the transcription termination site. Genome coverage using the rightmost base of each alignment was counted and normalized by counts-per-million using a customized Julia script using the XAM.jl package. Peaks were separately called for each replicate using Peaks.jl package with a width of 2. Peaks from the two replicates were merged with raw coverage values as peak heights. The heights of merged peaks were then normalized using the counts-per-million method. Classification of transcription termination sites (TTS) followed the procedure of ^27^. Primary transcription termination sites were defined as the peak with the highest coverage within 100 bp downstream of a gene (including the last base of the gene), while secondary TTSs were the other peaks in the region. TTS defined as internal were those between the start and end of a gene (excluding the last base). Antisense TTSs were on the opposite strand and between 50 bp upstream and downstream of a gene. TTSs that did not fall in above categories were classified as orphan TTSs. To identify potential Rho-independent terminator motifs, we used MEME ^28^ with the following parameters: -dna -oc . -nostatus -time 14400 -mod zoops -nmotifs 3 -minw 6 -maxw 50 -objfun classic - markov_order 0.

### Generation of chromosomal deletion mutants

*V. parahaemolyticus* deletion mutant strains were generated via homologous recombination & counterselection with the suicide plasmid pDM4, as described previously ^16^. Plasmids were generated by overlap PCR combined with restriction digests/ligation, or three fragment seamless cloning reactions (Sangon).

As an example, deletion of *lafK* is described. Approx. 1kb upstream and 1 kb downstream of the region to be deleted was amplified with primers DFO-0020/0021 and DFO-0022/0023 from WT genomic DNA ((gDNA, DFS-0191), respectively, which introduced a short (approx. 25 bp) region of homology between the two 1kb fragments for overlap PCR, as well as *Xba*I and *Xho*I restriction sites on the outside for cloning into pDM4. The upstream and downstream fragments were mixed in an equimolar ratio and subjected to overlap PCR using DFO-0020/0023, and pDM4 was amplified with DFO-0018/0019 and digested with *Dpn*I to remove template DNA. The overlap PCR product and pDM4 were digested with *Xba*I/*Xho*I and then ligated overnight with T4 DNA ligase. Ligations were transformed into DH5α λ*pir* and Cm^R^ colonies were verified by colony PCR with DFO-0044/0045. The verified plasmid (pSSvSH22) was chemically transformed into *E. coli* SM10 λ*pir or* MFD*pir,* followed by conjugation into the *V. parahaemolyticus* wild-type strain (DFS-0191). Transconjugants with single crossovers were first selected on MMM (minimal marine medium) plates with 10 µg/ml Cm. Single colonies were then transferred to MMM plates containing 15% sucrose to counter-select for double-crossover strains. Deletion clones were verified by colony PCR with DFO-0024/0023 and checked for Cm sensitivity. For deletions using MFD*pir*, clones were first selected on LB with 10 µg/ml Cm, followed by counterselection on LB + 20% sucrose. All deletion clones were tested for Cm sensitivity.

In some cases, a kanamycin resistance cassette was included via overlap PCR. As an example, deletion of *hfq* is described. The 1kb upstream region was amplified with DFO-0034/0224 and downstream region with DFO-0225/0037. A non-polar kanamycin resistance cassette was amplified from pKD4 ^29^ with DFO-0220/0221. The three fragments were joined by overlap PCR with DFO-0034/0037. Following digestion with *Xba*I/*Xho*I, the amplicon was ligated into similarly digested pDM4 (amplified with DFO-0018/0019) and transformed into *E. coli* DH5α λ*pir*. A positive clone was verified by colony PCR with DFO-0044-0045. Deletion mutants were generated as described above, except kanamycin was included in selection plates. Deletion mutants were verified with DFO-0038/0235.

### Plasmids for sRNA overexpression

To overexpress sRNAs using IPTG induction, we cloned them behind the Ptac promoter in conjugatable plasmid pMMB207 ^30^. For *vcrX,* the pMMB207 plasmid was amplified with DFO-0837/0565, and *vcrX* was amplified from wild-type gDNA using DFO-0838/0692. Vector and insert were joined by seamless cloning and transformed into *E. coli* DH5α. A positive clone (pSSvSH131) was verified by colony PCR with DFO-0306/0307 and Sanger sequencing with DFO-0306. For some flagellin gene/sRNA insertions in pMMB207, we amplified the vector with DFO-0437/0438 and insert, e.g., *flaY*, with DFO-1222/1223. Vector and insert were joined by seamless cloning and transformed into *E. coli* DH5α. A positive clone (pSSvSH194) was verified by colony PCR with DFO-0306/748 and Sanger sequencing with DFO-0306.

For induction using the pBAD promoter, we first modified pMMB207 to include *araC* and PBAD promoter sequences as follows. The pMMB207 vector was amplified with DFO-0287/0565, and *araC*-PBAD was amplified with DFO-0285/0564 from pBAD33. After digestion with *Xho*I, the vector and insert were ligated and transformed into *E. coli* DH5α. The plasmid was verified by colony PCR with DFO-0285/0307. We then inserted the sequence of the sRNA as follows, with FlaX provided as an example. The pZJ14 plasmid was amplified with DFO-0564/0565, and *flaX* was amplified from wild-type gDNA with DFO-0560/0561 to add regions complementary to pZJ14. Vector and insert were joined by seamless cloning and transformed into *E. coli* DH5α. A positive clone (pSSvSH87) was verified by colony PCR with DFO-0251/0307 and Sanger sequencing with DFO-0251.

Cloned sRNA sequences are listed in **Table S15**. In cases where a minor flagellin gene was inserted along with the downstream sRNA, the insert started at the TSS and ended at the sRNA terminator.

### Small RNA pulse-overexpression differential expression analysis

For data analysis, Fastp ^18^ version 1.0.1 was used for quality control of the raw reads with default parameters. Reads were then reverse complemented and mapped to the corresponding reference genome using Segemehl version 0.3.4 ^19^ with default parameters. FeatureCounts version 2.1.1 was used for gene expression quantification with parameters “-B -C -f -p -s 1 -O --fraction --countReadPairs --ignoreDup –primary”. Differential expression was calculated using R package DESeq2 ^21^ version 1.42.1 using the output from featureCounts ^20^ without normalization. Other general sequencing file and alignment file manipulations (e.g., read reverse complementation, alignment statistics, genome coverage file generations, etc.) and calculations (*e.g*., TPM normalization) were performed using Samtools ^22^ version 1.22.1 and Julia ^23^.

### Translational GFP fusions for sORF validation

A conjugatable sfGFP plasmid for translational fusions, pZH3, was first generated based on pMMB207. We first amplified sfGFP from its second codon from pXG10-SF ^9^ using DFO-0128/0139 and pMMB207 using DFO-0337/0185. After digestion with *Pst*I, the vector and insert were ligated and transformed into DH5α. A positive clone (pZH3), including the Ptac promoter fused to the second codon of sfGFP, was verified by colony PCR with DFO-0306/0748. This served as an empty control plasmid and template for sORF insertion. To insert *vcdP*, *spfP*, *napE*, *ryhP* to the 5th codon, we amplified pZH3 with DFO-0337 and a forward primer including the sORF (DFO-0341, 0288, and 0339, respectively). This included the TSS and ribosome binding site for *spfP*, and approximately 20 bp upstream of the start codon for *vcdP* and *ryhP*. Following phosphorylation with polynucleotide kinase (Thermo), inverse PCR products were ligated and transformed into *E. coli* DH5α. Positive clones (pZH5, pZH6, pZH7, pZH8) were verified by colony PCR with DFO-0306/0168 and Sanger sequencing with DFO-0306. Point mutations were introduced into putative start codons by inverse PCR with mutagenic primers, with oligonucleotides and resulting plasmids and sequences presented in **Tables S12** - **S14**. The rest of the *ryhP* sORF was added (to the penultimate codon) by inverse PCR on pZH5 with DFO-0444/0128, PNK treatment, ligation, transformation, and verification as described above to give pZH10, containing the full length ORF. To generate *V. cholerae* and *V. vulnificus* full-length *ryhP* fusions, we used strains E7946 (El Tor, Ogawa) and ATCC 27562, respectively, as template for PCR.

### Plasmids for measuring post-transcriptional regulation in *E. coli*

We used a previously described two-plasmid system in *E. coli* ^9^ for testing target regulation. As an example, we describe cloning for FlaX and its targets. First, the *flaX* sequence was cloned into pZE12 as follows. pZE12-luc was amplified by inverse PCR with DFO-0144/0145. The *flaX* sequence (424 nt; **Table S15**) was amplified from *V. parahaemolyticus* gDNA using DFO-0693/0694, adding sequences complementary to pZE12. Vector and insert were joined by seamless cloning and transformed into *E. coli* DH5α. A positive clone (pSSvSH113) was identified by colony PCR with DFO-0142/0143 and Sanger sequencing with DFO-0142. Point mutations in the *flaA* interaction site (binding site 3, **Fig. S7B**, M1) were introduced by inverse PCR with mutagenic primers DFO-0784/0785, followed by *Dpn*I digestion, DH5α transformation, and verification of the expected mutations (plasmid pSSvSH152; **Table S15**) by colony PCR and sequencing with the same primers as above. For targets, with the *flaA* 5’ UTR as an example, the region encompassing the TSS and first five codons was amplified with DFO-0695/0696 from *V. parahaemolyticus* gDNA, adding sequences complementary to pXG10-SF, and pXG10-SF was amplified with DFO-0157/0158. Vector and insert were joined by seamless cloning. Positive clones (pSSvSH108) in DH5α were validated by colony PCR with DFO-0167/0168 and Sanger sequencing with DFO-0167. Compensatory point mutations in the *flaA* 5’ UTR in binding site 3 (M1’) were introduced by inverse PCR with mutagenic primers DFO-0845/0846 as described above, and the resulting plasmid was verified by colony PCR and sequencing (pSSvSH145; **Table S15**). pZE12-sRNA plasmids were then introduced into *E. coli* DH5α carrying the appropriate pXG10-SF plasmid by electroporation.

### Transcriptional reporters in *V. parahaemolyticus*

GFP reporters for the *spf* and *vcrX* promoters were generated in a pMMB207-related plasmid carrying GFP with an unrelated RBS and no *lac* promoter as follows. Plasmid pSSvSH45 ^31^ was amplified by inverse PCR with primers DFO-0351/0180 and digested with *Dpn*I. The spf or *vcrX* promoters, including ∼200 bp upstream of the transcription start site, were amplified from RIMD 2210633 WT genomic DNA with CSO-1348/1339 and CSO-1349/1341, respectively. The promoter fragments were assembled with the pMMB207-RBS-*gfp* backbone by seamless cloning. Positive clones were verified by colony PCR with DFO-0167/0168 and sequencing with DFO-0168. Plasmids were introduced into the WT or crp strain (DFS-2386) using conjugation with *E. coli* MFD*pir*.

### Additional genomics analysis

The VN-5003-Wue strain was sequenced at Novogene Europe using the PacBio sequencing platform. The genome was assembled from raw subreads using Flye v2.9.2 ^32^ with option “--pacbio-raw”. To generate a phylogenetic tree of *Vibrio* and other species, a multiple sequence alignment of VP0014 (*gyrB*) and its homologs from the indicated strains and species was generated using Clustal Omega ^33^ (https://www.ebi.ac.uk/jdispatcher/msa/clustalo). The phylogenetic tree was then generated using IQ-TREE v3.0.1 ^34^ with the HKY model (-m HKY) and 100 bootstrappings (-b 100).

---

**Figure S1.**
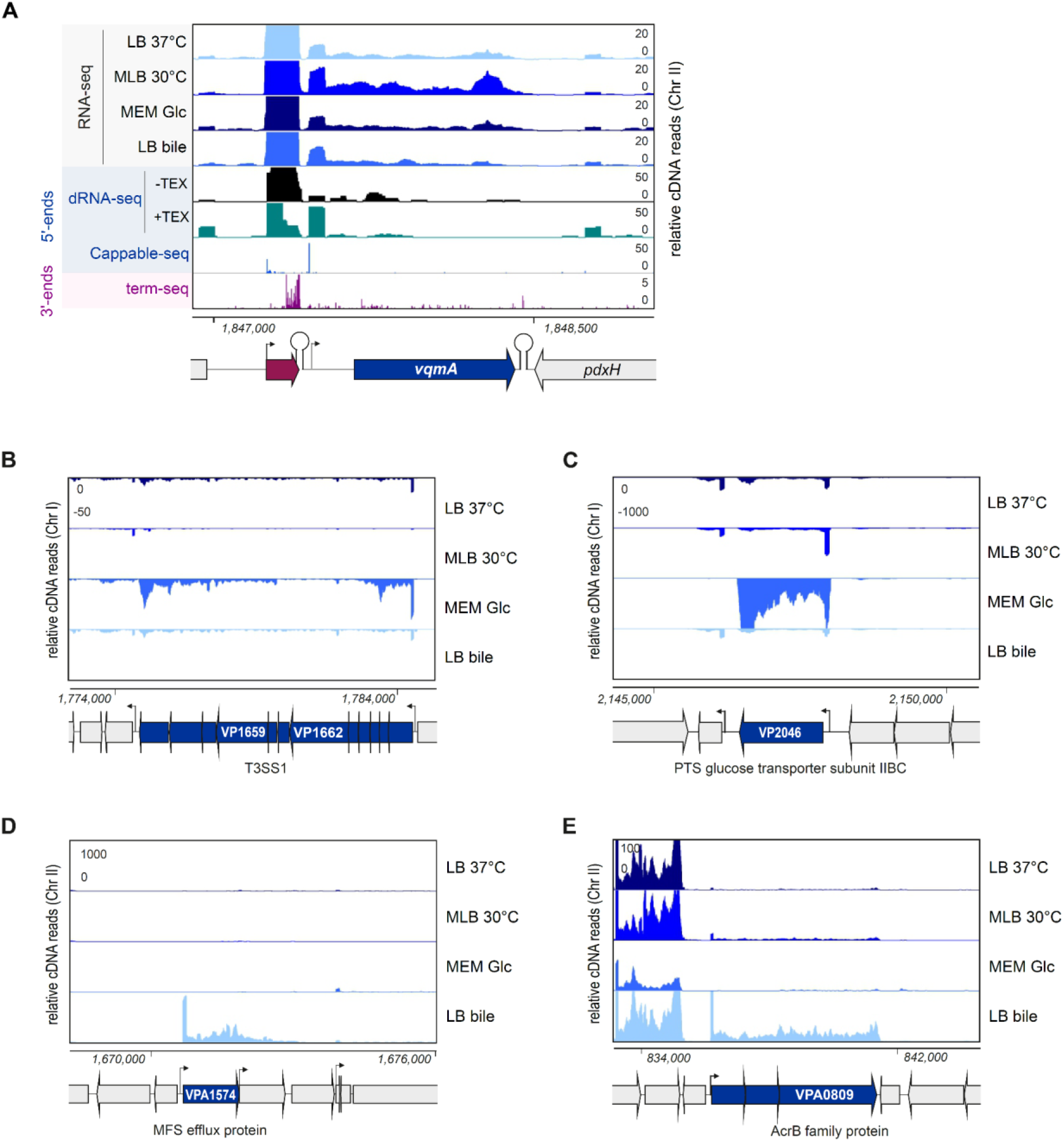
RNA-seq coverage. **(A)** Coverage at the *V. parahaemolyticus vqmRA* locus for RNA-seq libraries generated in this study. **(B & C)** RNA-seq coverage under four conditions for example genes induced in the T3SS1-inducing condition (MEM + Glc). **(D & E)** RNA-seq coverage under four conditions for example genes induced in the T3SS2-inducing condition (LB + bile salts). Bent arrows: TSS.

**Figure S2.**
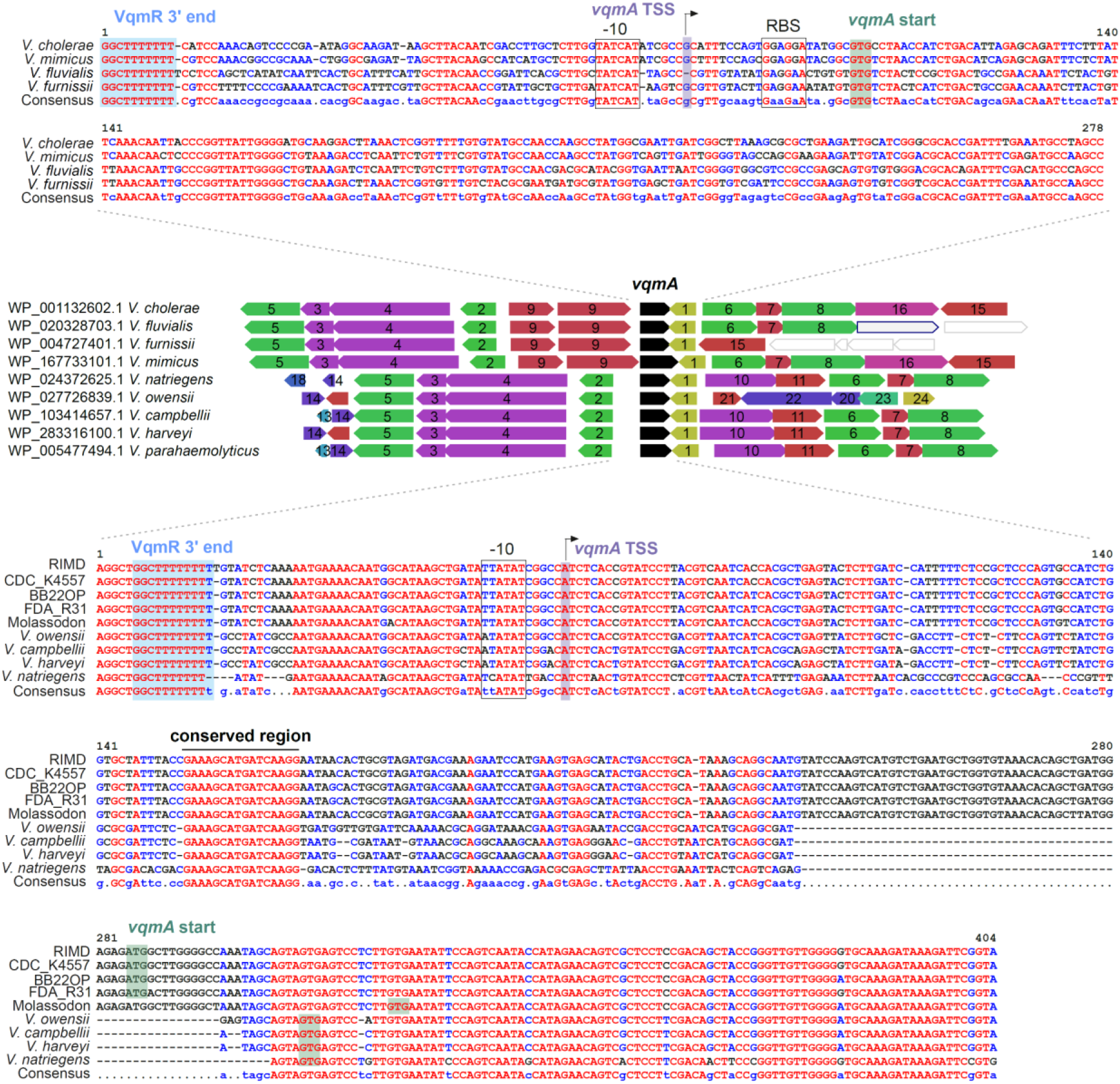
The *Vibrio vqmR-vqmA* locus. (*Middle*) Synteny analysis based on the *V. parahaemolyticus* VqmA sequence (black) using webflags ^1^. Numbers: homologous genes. Intergenic regions are to scale. (*Top & bottom*) Alignment of two types of *vqmA* region from *Vibrio* species. Blue: end of *vqmR*. Purple/bent arrow: TSS for *vqmA* based on *V. parahaemolyticus* (this study) *or V. cholerae* ^2^ dRNA-seq. Green: annotated *vqmA* start codons, except for *V. cholerae*, which is independently validated ^2^.

**Figure S3.**
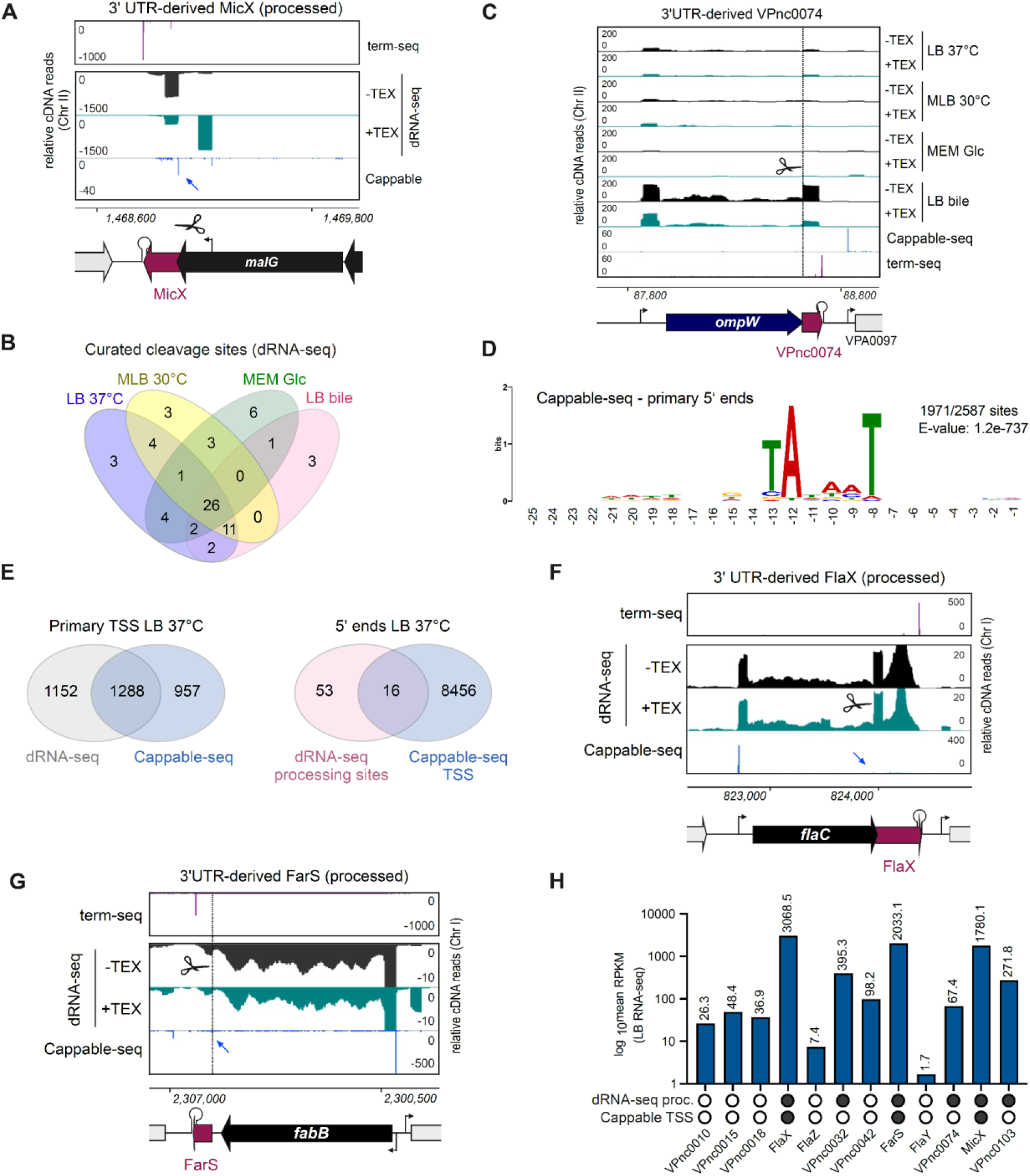
Detection of 5’ ends by dRNA-seq and Cappable-seq. **(A, C, F, G)** cDNA library coverage in *V. parahaemolyticus* for the 3’ UTR-derived sRNAs MicX, FlaX, and FarS. Blue arrows: Cappable-seq peaks. Scissors/vertical dashed line: dRNA-seq processing sites. **(B)** Curated processing sites based on dRNA-seq. **(D)** MEME motif analysis of the 50 bp upstream of primary TSS identified by Cappable-seq in LB. **(E)** Comparison of primary TSS identified by dRNA-seq (left) or processing sites identified by dRNA-seq (right) and Cappable-seq TSS. For dRNA-seq, only sites from the LB 37°C condition were used. **(H)** Comparison of 3’ UTR-derived sRNA expression in LB and detection of a dRNA-seq processing site or Cappable-seq TSS in the LB 37°C condition.

**Figure S4.**
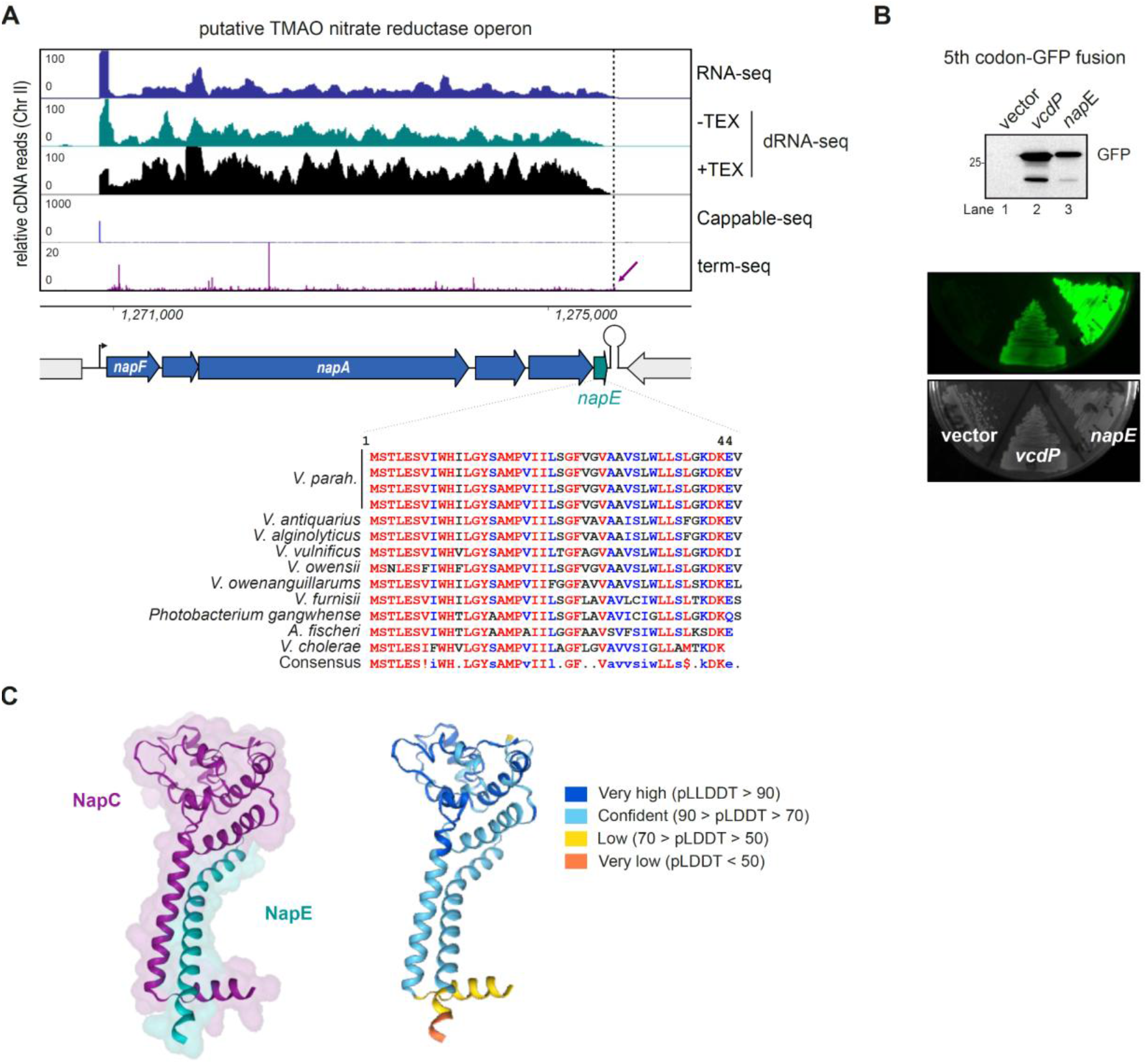
Putative small NapE-like component of *Vibrio* nitrate reductase. **(A)** (*Top*) cDNA library coverage in *V. parahaemolyticus* at the TMAO (trimethylamine *N*-oxide) nitrate reductase operon. (*Bottom*) protein sequence alignment for putative *napE* homologs identified by tblastn with the *V. parahaemolyticus* protein sequence. Sequences were aligned with multalin ^3^. Dashed vertical line: 3’ end of RNA-seq coverage. Purple arrow: term-seq peak associated with end of RN-seq coverage for the operon. **(B)** Western blot (*top*) and fluorescence (*bottom*) for a *napE* 5th codon translational GFP fusion. The sORF *vcdP* and insertless GFP vector served as positive and negative controls, respectively. **(C)** Alphafold multimer ^4^ prediction for NapC (VPA1201) and NapE. The prediction was performed at Colabfold ^5^ and visualized with PAE Viewer ^6^.

**Figure S5.**
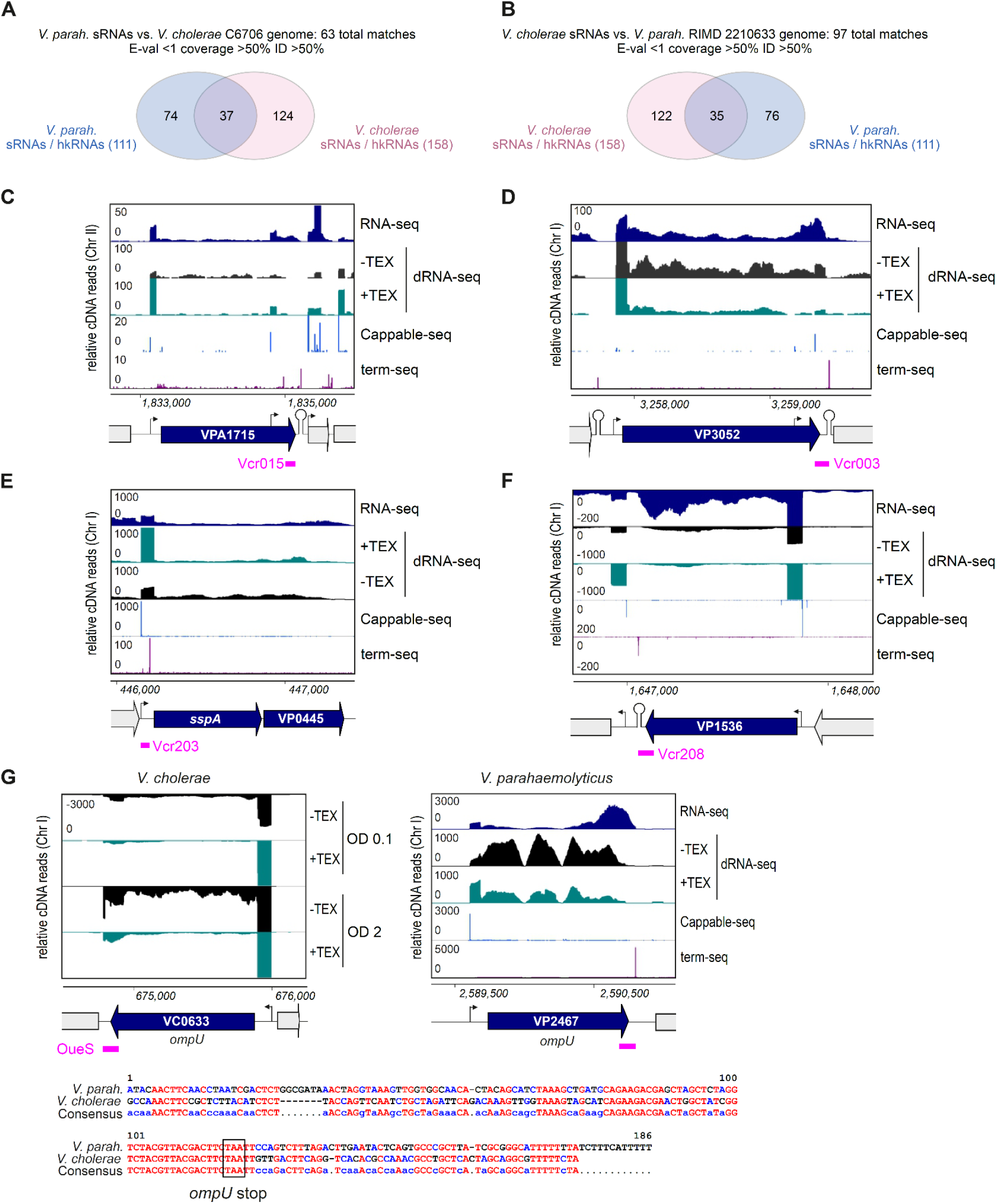
*V. cholerae* and *V. parahaemolyticus* sRNA overlap. **(A)** Number of experimentally-identified *V. parahaemolyticus* sRNAs (n=111) with BLASTn matches that overlap with experimentally-identified *V. cholerae* C6706 sRNAs ^2,7^ (n=158). hkRNA: housekeeping RNA. **(B)** Number of *V. cholerae* sRNAs with significant BLASTn matches in the *V. parahaemolyticus* RIMD 2210633 genome. **(C-F)** Example cDNA library coverage for regions with significant homology to *V. cholerae* sRNAs that were not identified as sRNAs by ANNOgesic using our dataset. **(G)** cDNA library coverage at the *ompU* porin region for *V. cholerae*, encoding OueS, and the cognate region in *V. parahaemolyticus*. Regions with homology identified by BLAST are indicated with pink bars. See also **Table S8** for all matches with BLAST outputs. *V. cholerae* sRNA-seq data is from ^2^.

**Figure S6.**
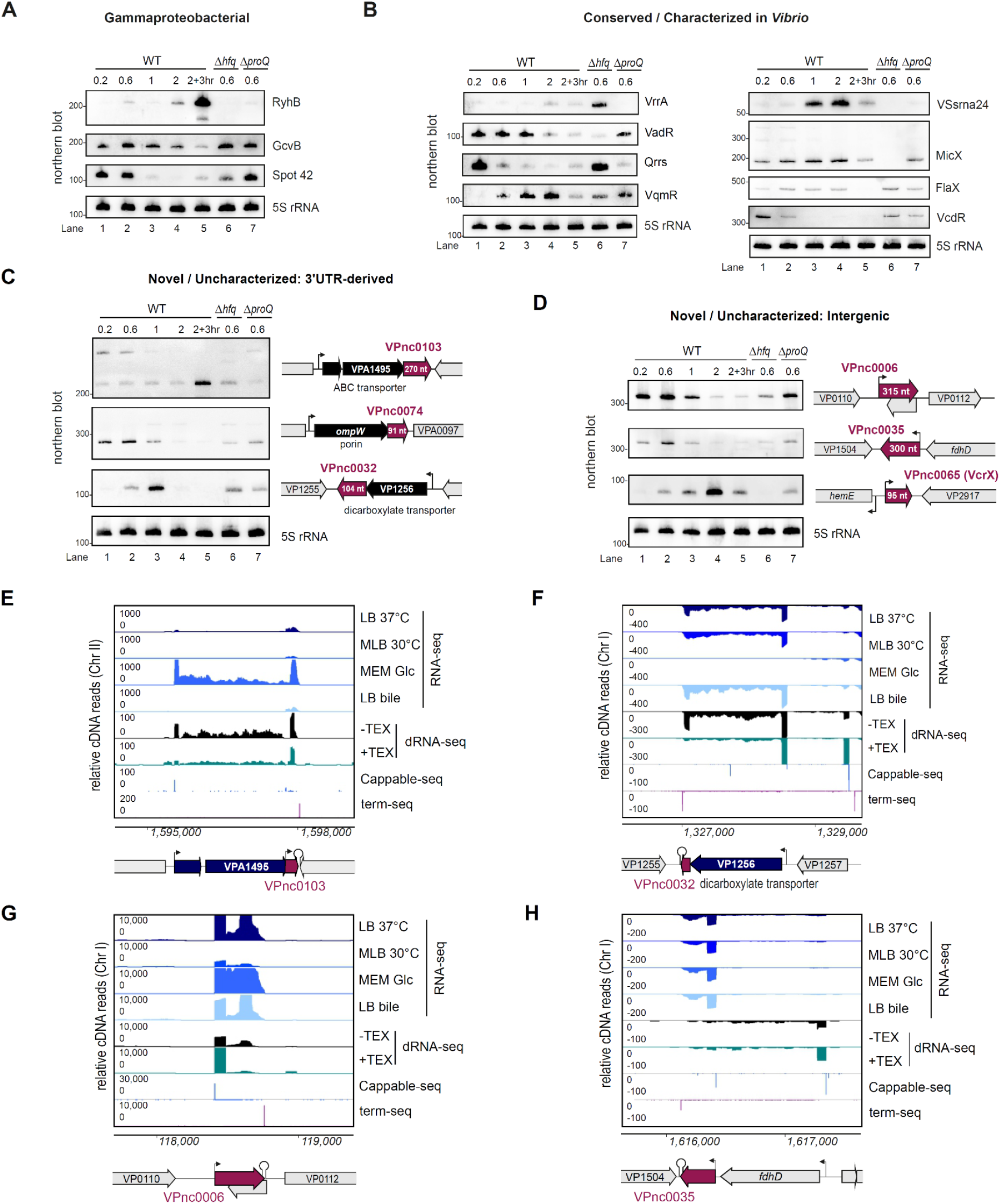
Expression of conserved, taxon-specific, and new sRNAs. **(A-D)** Northern blot validation of sRNAs in different growth phases in rich medium. RIMD 2210633 WT or isogenic Δ*hfq*/Δ*proQ* mutants were grown in MLB (LB + 3% NaCl) to the indicated OD_600_ at 30°C. Total RNA was analysed by northern blot using biotin-labeled probes (**Table S12**). 5S rRNA: loading control. Genomic context cartoons are not to scale. Representative of two independent cultures. **(E-H)** cDNA library coverage for novel/uncharacterized sRNAs validated by northern blot. Panels E & F: 3’ UTR-derived sRNAs (see main Fig. 3C). Panels G & H: intergenic sRNAs (related to main Fig. 3D). Bent arrows: transcription start sites.

**Figure S7.**
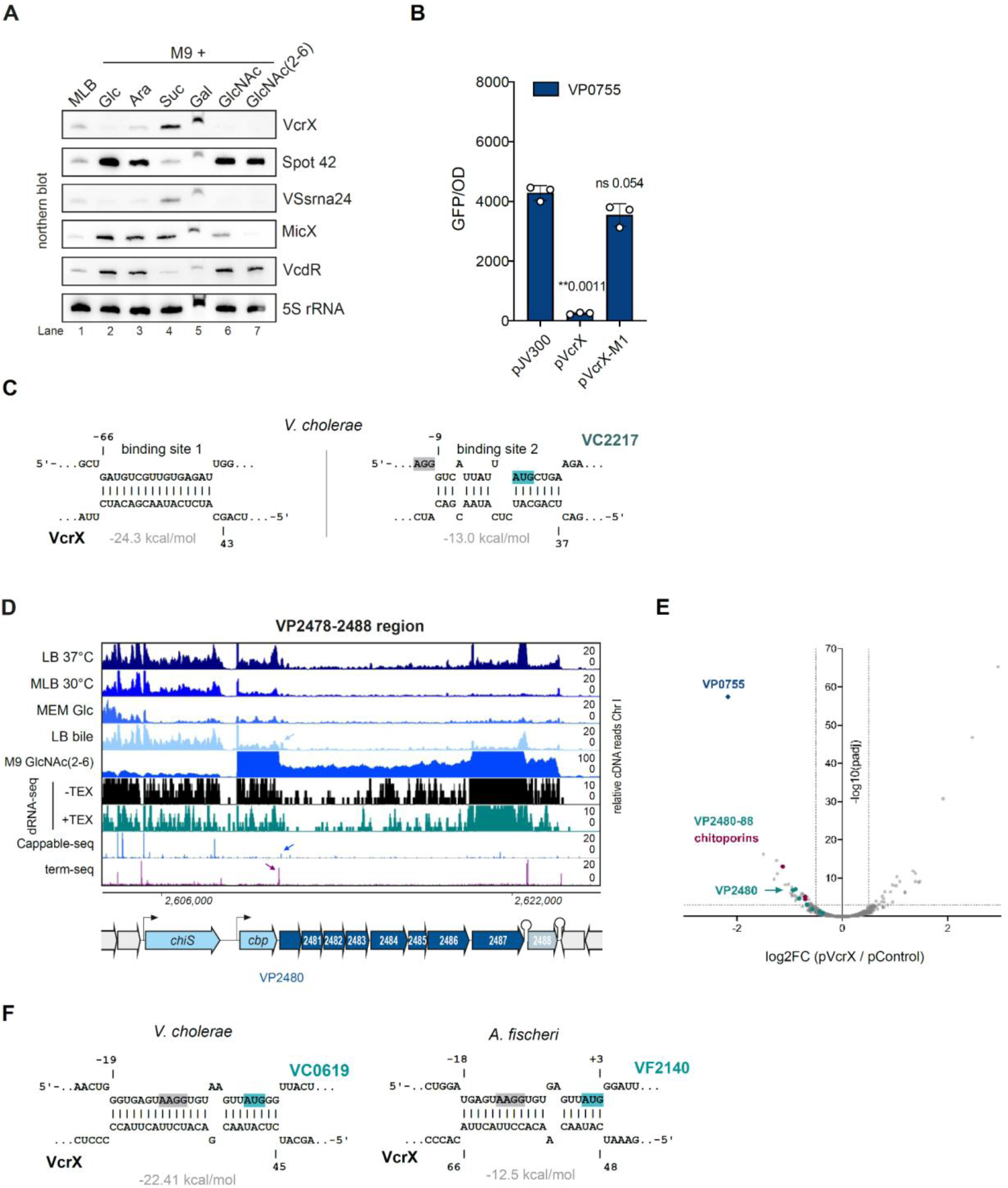
VcrX expression and targets. **(A)** Northern blot analysis of *crp* mRNA and sRNA expression in M9 supplemented with different carbon sources at 0.2%. MLB: rich medium. Glc: glucose; Ara: arabinose; Suc: succinate; Gal: galactose; GlcNAc: *N*-acetylglucosamine; GlcNAc(2-6): 2-6 unit chito-oligosaccharides. 5S rRNA: loading control. **(B)** VP0755 translational fusion expression in presence/absence of VcrX. n=3, Student’s t-test. **p<0.01, ns: not significant. **(C)** Predicted base pairing between *V. cholerae* VcrX and VP0755 homolog VC2217. **(D)** Coverage at the VP2478-VP2488 region. Blue arrow: Cappable-seq peaks 20 bp into VP2480. **(E)** RNA-seq after VcrX pulse-overexpression in M9 + GlcNAc(2-6). n=2. Dashed lines: log2FC = |0.5|; - log10(p) = 3. **(F)** Predicted base pairing between *V. cholerae/A. fischeri* VcrX and VP2480 homologs VC0619/VF2140.

**Figure S8.**
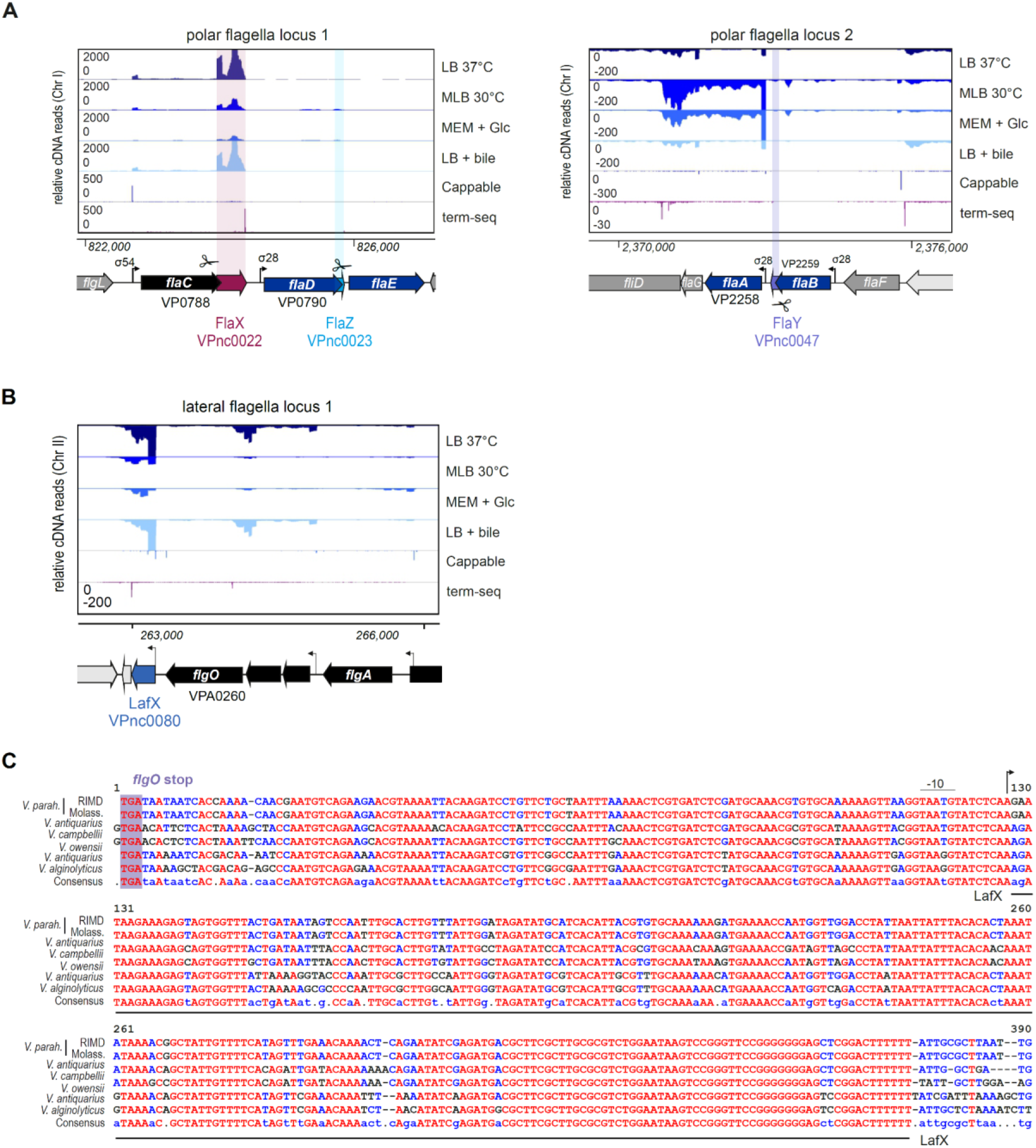
Small RNAs encoded in flagellar regions. **(A)** cDNA library coverage for *V. parahaemolyticus* FlaX, FlaY, and FlaZ sRNAs encoded in the 3’ UTR of polar flagellin genes. **(B)** cDNA library coverage for *V. parahaemolyticus* LafX, encoded adjacent to the first lateral flagella gene cluster on Chromosome 2. **(C)** Alignment of LafX homologs encoded adjacent to *flgO* identified by BLAST in *Vibrio* species.

**Figure S9.**
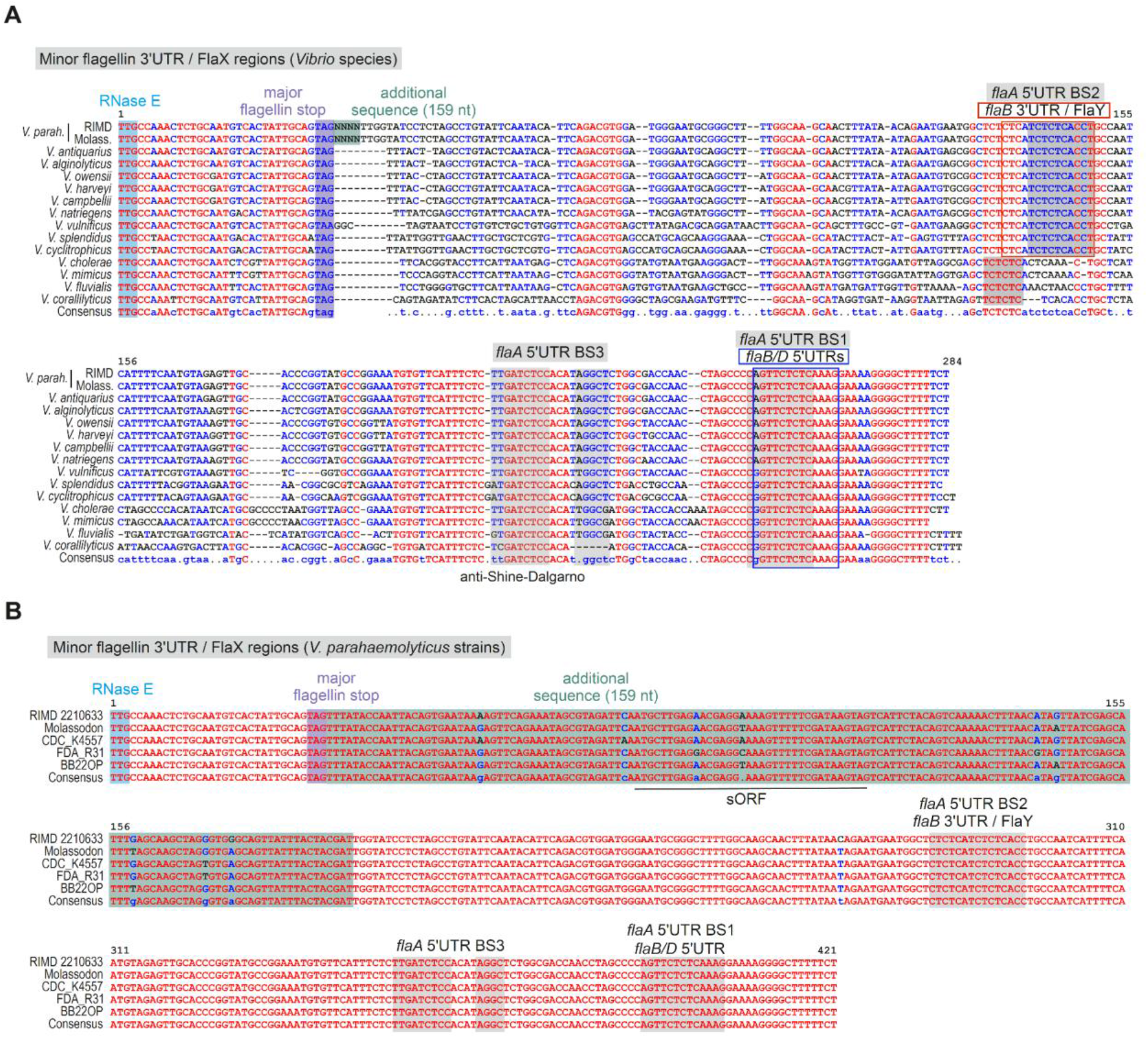
Sequence analysis of polar flagellins and FlaX. **(A)** Sequence alignment of FlaX regions from *Vibrio* species. Regions downstream of the major flagellin identified via homology and genomic location compared to *V. parahaemolyticus* and *V. cholerae* were included. Note that in *V. parahaemolyticus* and several other species, the major flagellin is termed “FlaC”, while in others it is unnamed or called “FlaA” (*e.g.*, *V. cholerae*, see also panel A). Green/NNN: inserted sequence in *V. parahaemolyticus* strains (see panel C). Grey: base-pairing regions with the indicated flagellin mRNAs in *V. parahaemolyticus*. BS: binding site. **(B)** Sequence alignment of FlaX regions from *V. parahaemolyticus* strains. Green: inserted sequence. BS: binding/base-pairing site. Sequences were aligned with multalin ^3^.

**Figure S10.**
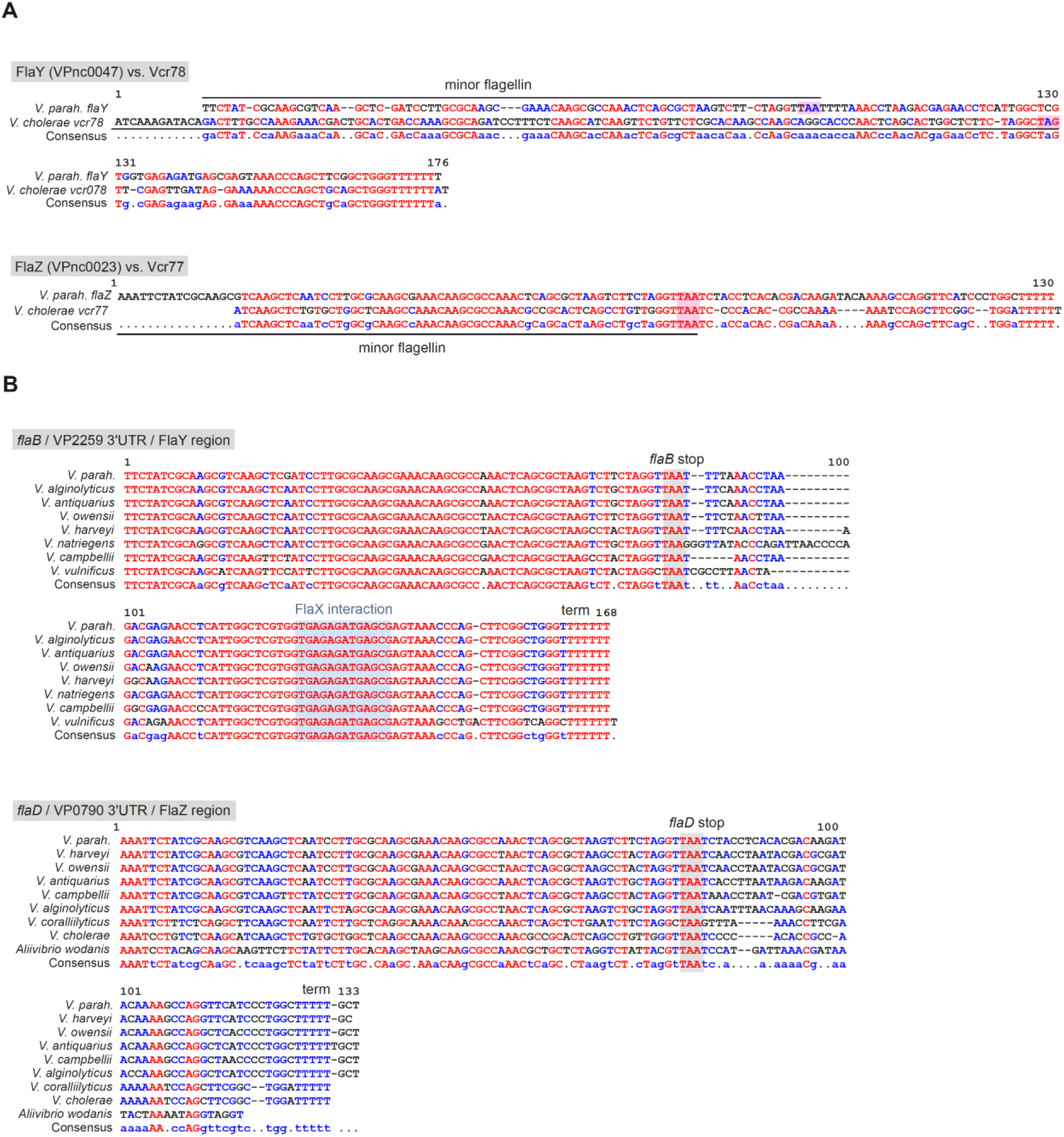
Conservation and sequence analysis of flagellin-associated sRNAs. **(A)** Alignment of *V. parahaemolyticus flaY/flaZ* and potential *V. cholerae* homologs Vcr078/Vcr077. **(B)** Alignment of VP2259/*flaB* homolog 3’ UTRs (*flaY*) or VP0790/*flaD* homolog 3’ UTRs (*flaZ).* Selected BLASTn matches from *Vibrio* species with coverage >90% and percent identity >80% were included. Sequences were aligned with multalin ^3^.

**Figure S11.**
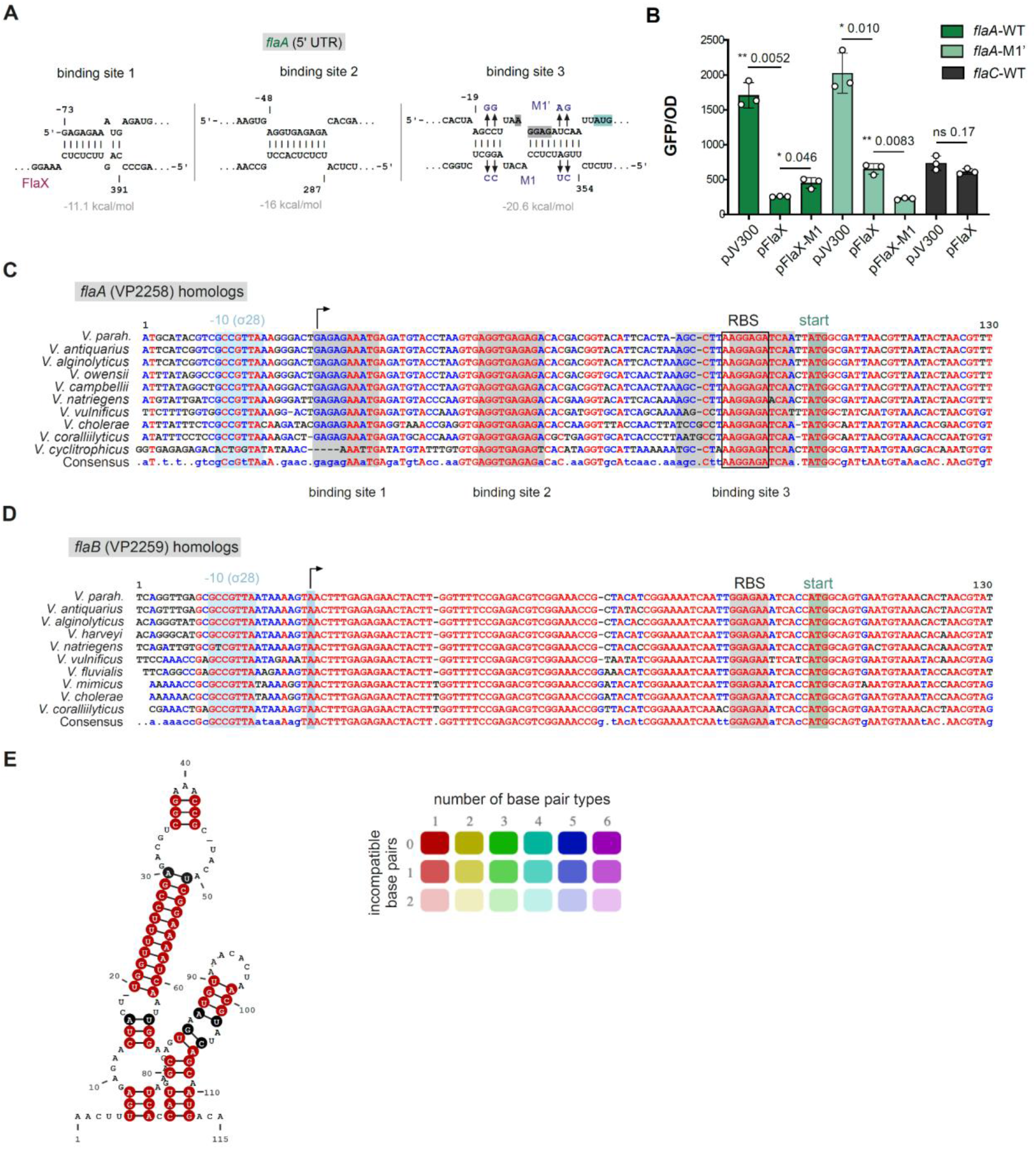
Sequence analysis of minor flagellin 5’ UTRs. **(A)** Predicted interactions between FlaX and the *flaA* 5’UTR. Binding sites are named according to ^8^. mRNA positions: relative to start codons. **(B)** FlaX effect on *flaA* and *flaC* 5’UTR translational reporters in an *E. coli* two-plasmid system ^9^. M1/M1’: see panel A. **(C & D)** Alignment of *flaA* (VP2258) or *flaB* (VP2259) promoter / 5’ UTR / first codons regions from *Vibrio* species. Bent arrows: TSS. Sequences were aligned using multalin ^3^. **(E)** Structural conservation of *flaB* 5’ UTRs using aligned sequences from panel C with RNAalifold ^10^.

**Figure S12.**
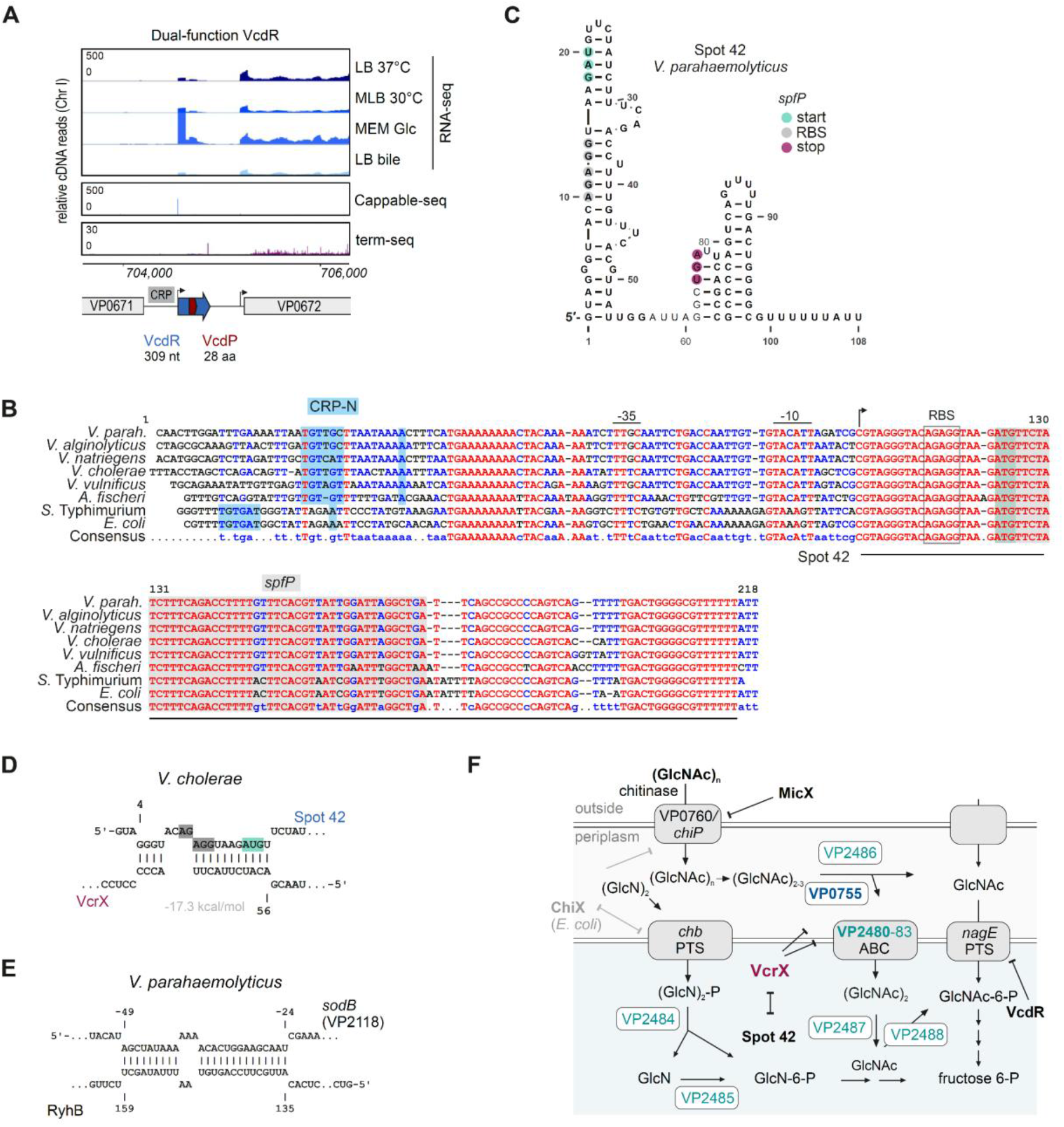
Dual-function sRNAs in *V. parahaemolyticus*. **(A)** RNA-seq coverage in *V. parahaemolyticus* for VcdR. Blue: sRNA. Red: sORF. **(C)** Alignment of Spot 42 RNA sequences and upstream promoter regions from Vibrionaceae and enterobacteria. Grey: *spfP* sORF. Boxed residues: putative ribosome binding site. CRP sites in *E. coli* and *Salmonella* homologs are from ^11^. CRP-N: based on ^12^ (*E. coli*) and homology to predictions in ^13^. The alignment was generated with multalin ^3^. **(C)** Predicted secondary structure of *V. parahaemolyticus* Spot 42 using RNAFold ^10^. Features of the *spfP* sORF are indicated. **(D)** Predicted interaction (IntaRNA) ^14^ between VcrX and Spot 42 from *V. cholerae*. **(E)** Predicted base-pairing between RyhB and *sodB* mRNA using IntaRNA ^15^. mRNA positions are relative to the start codon, while sRNA positions start from the TSS. **(F)** Expanded model of sRNAs involved in chitin regulation in *V. parahaemolyticus*.

**Table S15.**
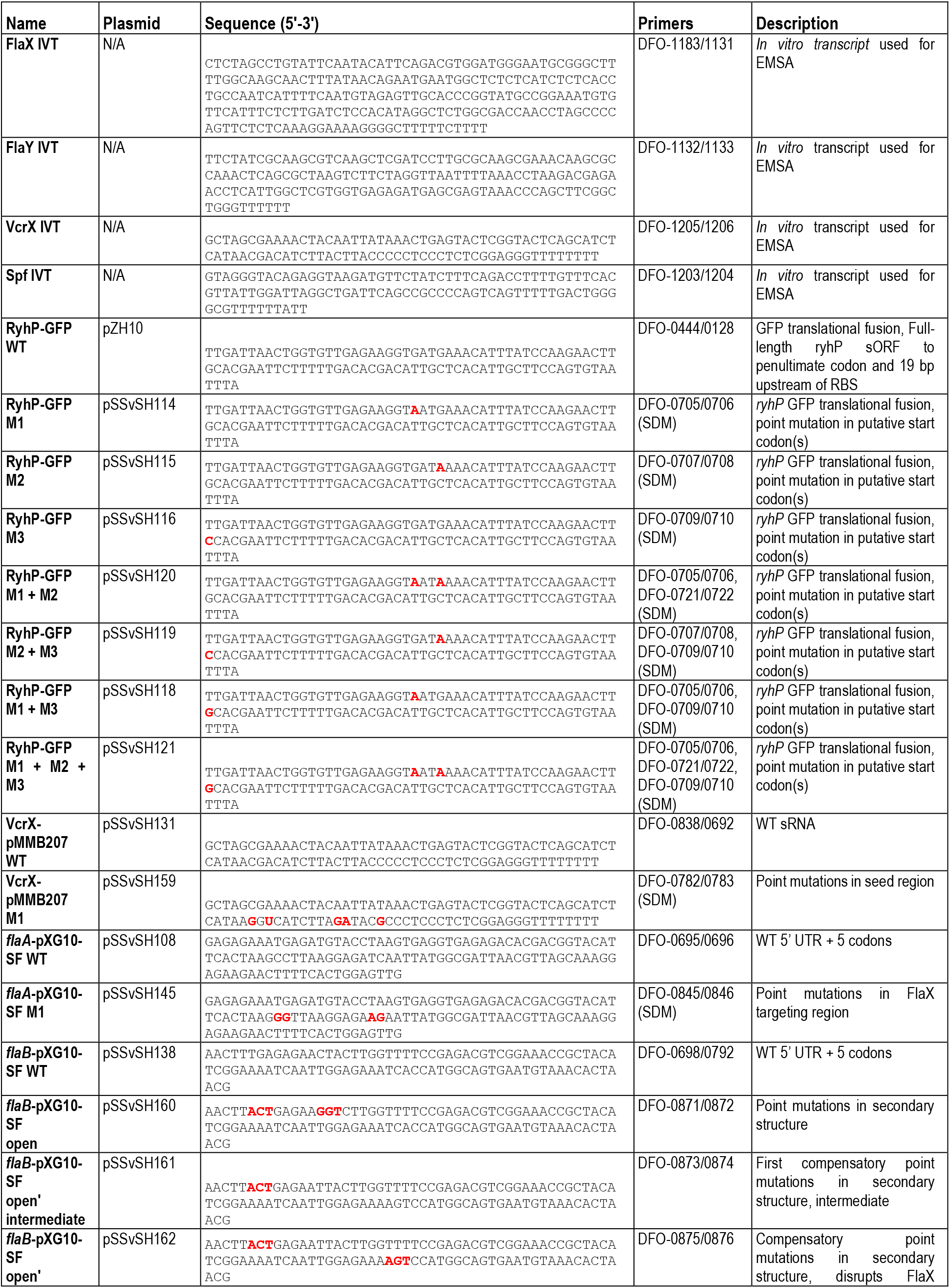
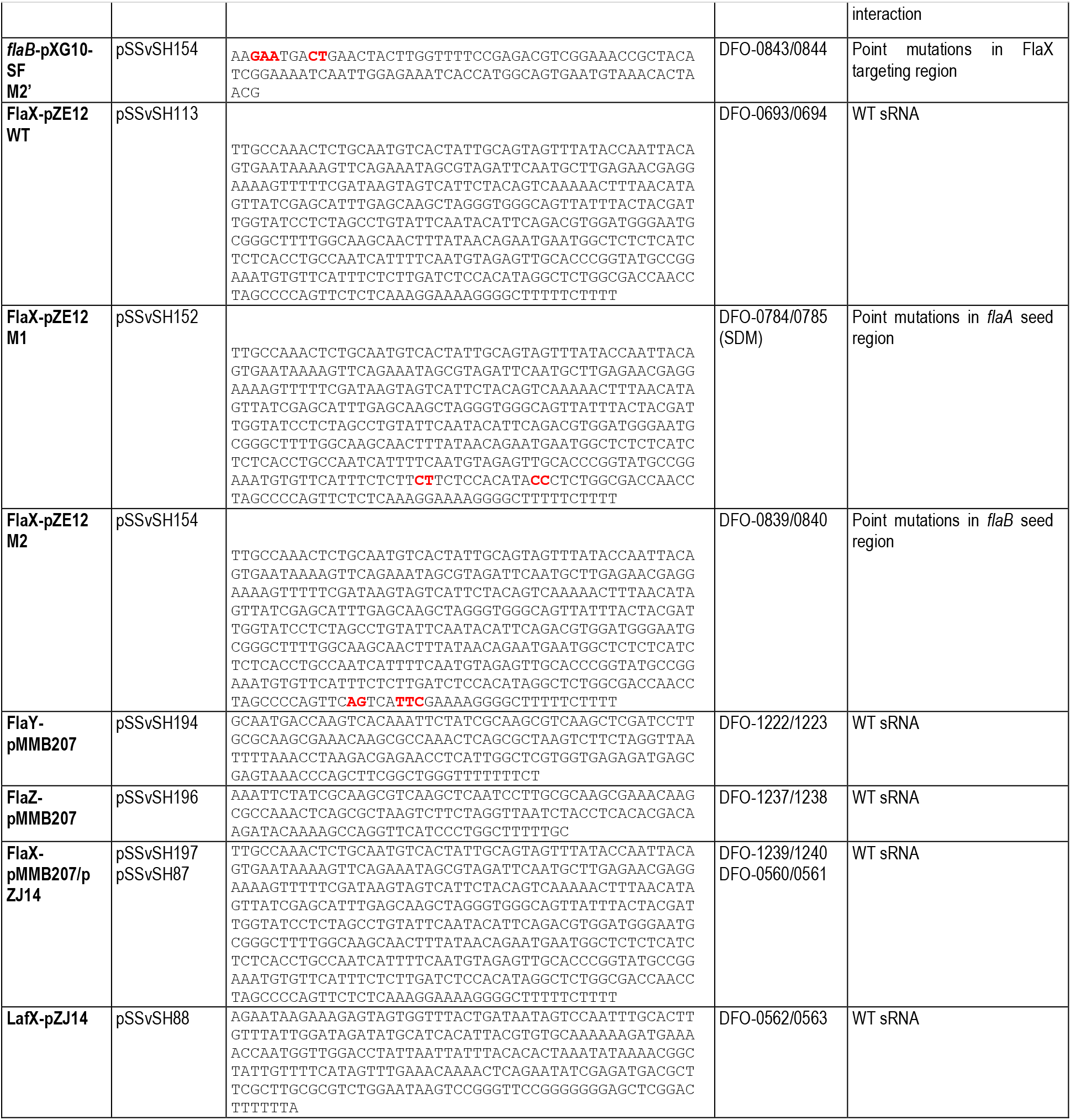
Cloned, *in vitro* transcript (IVT), and point mutant sequences. EMSA: electrophoretic mobility shift assay. SDM: site-directed mutagenesis. RBS: ribosome binding site. Mutated bases are indicated in red font.

